# Integration of Imaging-based and Sequencing-based Spatial Omics Mapping on the Same Tissue Section via DBiTplus

**DOI:** 10.1101/2024.11.07.622523

**Authors:** Archibald Enninful, Zhaojun Zhang, Dmytro Klymyshyn, Hailing Zong, Zhiliang Bai, Negin Farzad, Graham Su, Alev Baysoy, Jungmin Nam, Mingyu Yang, Yao Lu, Nancy R. Zhang, Oliver Braubach, Mina L. Xu, Zongming Ma, Rong Fan

## Abstract

Spatially mapping the transcriptome and proteome in the same tissue section can significantly advance our understanding of heterogeneous cellular processes and connect cell type to function. Here, we present Deterministic Barcoding in Tissue sequencing plus (DBiTplus), an integrative multi-modality spatial omics approach that combines sequencing-based spatial transcriptomics and image-based spatial protein profiling on the same tissue section to enable both single-cell resolution cell typing and genome-scale interrogation of biological pathways. DBiTplus begins with *in situ* reverse transcription for cDNA synthesis, microfluidic delivery of DNA oligos for spatial barcoding, retrieval of barcoded cDNA using RNaseH, an enzyme that selectively degrades RNA in an RNA-DNA hybrid, preserving the intact tissue section for high-plex protein imaging with CODEX. We developed computational pipelines to register data from two distinct modalities. Performing both DBiT-seq and CODEX on the same tissue slide enables accurate cell typing in each spatial transcriptome spot and subsequently image-guided decomposition to generate single-cell resolved spatial transcriptome atlases. DBiTplus was applied to mouse embryos with limited protein markers but still demonstrated excellent integration for single-cell transcriptome decomposition, to normal human lymph nodes with high-plex protein profiling to yield a single-cell spatial transcriptome map, and to human lymphoma FFPE tissue to explore the mechanisms of lymphomagenesis and progression. DBiTplusCODEX is a unified workflow including integrative experimental procedure and computational innovation for spatially resolved single-cell atlasing and exploration of biological pathways cell-by-cell at genome-scale.

## INTRODUCTION

The advent of high-throughput single-cell technologies has revolutionized our understanding of biological systems, enabling comprehensive analyses at the molecular level across diverse biological contexts^1-3^. These approaches, however, often overlook the spatial context within which molecular interactions occur, thus limiting our understanding of inter-cellular interactions occurring within tissues and organs^4^. Recent computational approaches have been developed to infer spatial information from single-cell RNA sequencing datasets^5,6^. In contrast, spatial omics was developed to address this limitation of missing spatial context using a wide range of approaches^7^. Array-based spatial transcriptomics such as Spatial Transcriptomics^8^ and Slide-seq^9^ utilize spatially barcoded surfaces to capture mRNA transcripts. Microfluidic-based approaches such as DBiT-seq^10^, spatial-ATAC-seq^11,12^, spatial CITE-seq^13^, and spatial-CUT&Tag^14^ employ microfluidics chips to deliver spatial barcodes to profile the transcriptome, proteome, or epigenome. Imaging-based methods such as MERFISH^15^ and seqFISH^16^ are multiplexed versions of single-molecule FISH (smFISH)^17^ which utilize combinatorial labeling and sequential imaging to achieve subcellular visualization and quantification of transcripts. Similarly, in situ sequencing technologies such as STARmap^18^, ISS^19^, and FISSEQ^20^ sequence nucleic acids directly within preserved tissues or cells, retaining their spatial context. This is typically achieved by incorporating sequencing-by-synthesis or sequencing-by-ligation methods combined with imaging to visualize and map gene expression patterns at high spatial resolution. Broadly, in situ sequencing or imaging techniques provide higher spatial resolution and sensitivity but have limited genome coverage for unbiased exploration of biological mechanisms. As such, carefully curated gene panels must be designed which can limit its discovery potential. On the other hand, sequencing-based methods have the drawback of lower resolution but tend to be transcriptome-wide. Techniques for profiling the proteome through either antibodies labeled with DNA barcodes visualized through cyclic hybridization and imaging of fluorescent readout sequences (CODEX)^21^ or iterative indirect immunofluorescence imaging (4i)^22^ have also been developed, offering single-cell resolution profiling of protein markers for in situ cell typing to generate spatial cell type maps.

We posit that the integration of imaging-based and sequencing-based spatial omics techniques brings the best of both worlds to yield a spatial cell atlas amenable for single-cell resolution cell type annotation and genome-wide mechanistic inquiry, providing a multidimensional view of molecular processes, maintaining the spatial context of biological interactions and enabling the understanding of complex cellular architectures and tissue microenvironments^23^. Current strategies for most spatial multi-omic technologies involve running single spatial omics assays separately on adjacent or serial tissue sections followed by computational data integration of the multimodal datasets. This has been applied to the study of human myocardial infarction^24^, the discovery of cellular niches in the human heart^25^ and the dissecting the tumor microenvironment^26,27^. Due to sample heterogeneity in cellular composition and architecture, even between adjacent tissue sections from the same block which differ in the exact cell types and orientation, integrating multimodal data computationally is suboptimal as perfect concordance between tissue sections is almost unattainable. As such, new methods for spatially resolved multi-modality and multi-omics measurements on the same tissue sections are highly desired. Recently, MiP-seq was developed to efficiently detect multiplexed DNAs, RNAs, and proteins in brain tissue at subcellular resolution^28^. Single-cell spatial multimodal metabolomics approaches such as scSpaMet were reported to combine protein-metabolite measurements of single immune and cancer cells^29^. While it would be ideal to integrate subcellular resolution image-based cell type identification with genome-wide spatial omics sequencing as well as computational registration for downstream analysis, the best solutions are yet to be established.

Here, we developed **D**eterministic **B**arcoding in **T**issue **s**equencing **p**lus (abbreviated as DBiTplus) which combines spatially resolved omics sequencing with highly multiplexed immunofluorescence imaging (CODEX) for the co-profiling of whole transcriptome and a panel of ∼50 protein markers on the same tissue section. We tested several methods to retrieve cDNA without damaging the tissue section but chose to use and optimize an enzymatic approach using RNaseH for the retrieval of cDNAs from tissue sections after the spatial barcoding workflow while maintaining tissue integrity and morphology prior to the CODEX workflow. A novel computational approach modified from the MaxFuse algorithm^30^ was developed to integrate the multi-omic spatial transcriptomic and CODEX datasets and the RCTD-like approach^31^ for DBiTplus spot cell type deconvolution and spot splitting to generate pure cell-type sub-spots. We applied DBiTplus to elucidate the process of embryogenesis in OCT-frozen and paraffin-embedded C57 mouse embryo sections. Applying DBiTplus to healthy human lymph node tissue and diseased lymphoma tissues, we demonstrated the capability to generate high-quality transcriptome and protein data from the same tissue section, overcoming the hurdles associated with data integration and registration from adjacent sections that show numerous slight differences in spatial composition.

## RESULTS

### Design and overview of DBiTplus

In the standard DBiT-seq workflow, mRNAs are reverse-transcribed in situ in the tissue matrix, and a microfluidic chip with 50 parallel channels delivers DNA barcodes A (Ai; i=1–50) and B (Bj; j=1–50) perpendicularly. The barcodes ligate to form a unique 2D array of barcoded spots. After tissue lysis, barcoded cDNAs are recovered, purified, and amplified for paired-end sequencing to generate a spatial gene expression map. This technique has been adapted to profile both the transcriptome, epigenome, proteome and more recently, applied to archival formalin-fixed paraffin-embedded tissue blocks^10-14,32,33^. Additionally, multiplexed versions of this technique have also been developed to profile up to nine unique tissue sections in parallel on the same tissue slide^34^. Central to the efforts to combine multiplexed immunofluorescence with DBiT-seq was to develop novel techniques to release cDNA from tissue sections while maintaining tissue integrity. Two chemical approaches (using NaOH and DMSO) and an enzymatic approach (using RNaseH) were tested (**Extended Data Fig. 1a**). NaOH and DMSO are known denaturing agents of DNA^35^. Following successful retrieval of the cDNA and preparation of a sequencing library, the intact tissue section is imaged via CODEX, and routine H&E staining is performed on the same tissue section (**Fig. 1a**). For FFPE samples, the Patho-DBiT workflow^33^ is used until the spatial barcoding process is completed following which the tissue section is incubated at 55°C with a mix of Triton X-100 and Thermostable RNaseH enzyme to break down RNA strands in RNA–DNA hybrids and to facilitate the diffusion through the permeabilized cell membranes. Another overnight incubation at 37°C is performed to increase the retrieval of cDNA from the tissue. Tubes with retrieved cDNA are pooled together and purified, template switched, amplified, and a sequencing library built. The intact tissue section can be stored at -20°C until CODEX staining. The intact FFPE tissue section undergoes brief rehydration and antigen retrieval steps (these are performed in the Patho-DBiT workflow) prior to staining with a panel of markers and imaging. After successful imaging, the flow cell is removed from the tissue slide (**See Methods**) and routine H&E staining is performed on the same tissue section. Similarly, for fresh frozen (FF) samples, the same DBiTplus workflow is followed as described minus steps specific to FFPE samples.

**Fig. 1.**
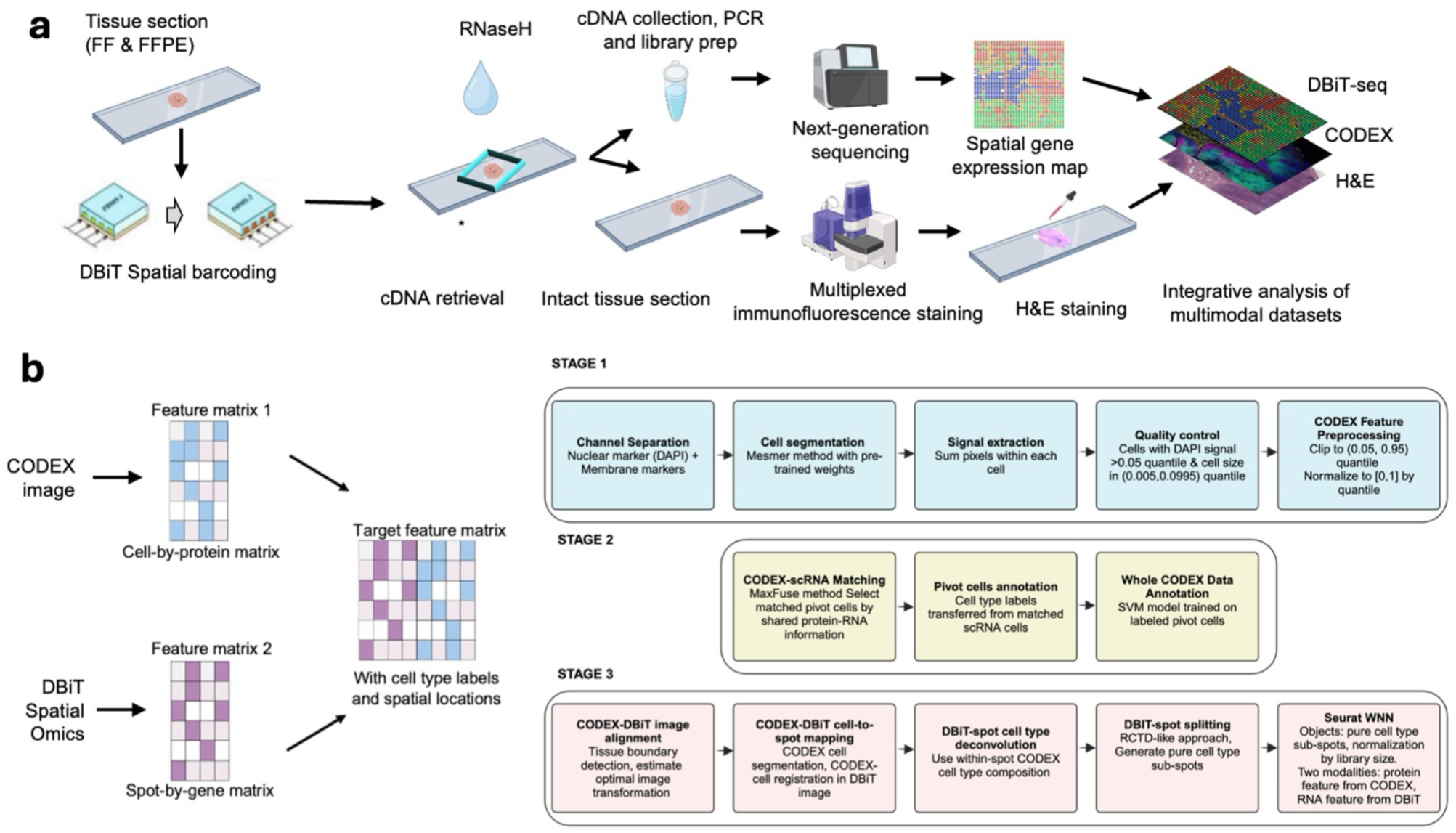
DBiTplus workflow and summary of bioinformatic workflow for integrative multimodal data analysis. **a** Workflow of DBiTplus technology. Both FF and FFPE tissue sections can be used in this workflow. Following reverse transcription and spatial barcoding, the tissue section is incubated with thermostable RNaseH for cDNA retrieval. The tubes are pooled, followed by purification of cDNA, PCR amplification and library preparation for sequencing. The intact tissue section is used for multiplexed immunofluorescence staining using the PhenoCycler Fusion system. Histological stain (H&E) is performed on the tissue following multiplexed immunofluorescence staining and integrative analysis of the multimodal datasets is performed. This approach demonstrates the ability to generate spatial transcriptomic, proteomic and histological data from the same intact tissue section. **b** A cell-by-protein matrix from the CODEX dataset and a spot-by-RNA matrix from the DBiTplus workflow are integrated to generate a long feature vector with the cell’s CODEX reads and the average RNA expression levels of all cells within the same DBiTplus spot that is of the same cell type. Detailed three-stage workflow for the integration of protein and RNA datasets. Cell segmentation using a Mesmer-based model followed by signal extraction, quality control and CODEX feature processing. MaxFuse algorithm is utilized to match pivot cells shared between protein and single-cell RNA seq reference dataset, followed by the annotation of pivot cells and the whole CODEX dataset. For tumor tissue, tumor cells were identified by CODEX markers first. MaxFuse was then applied on non-tumor cells. The CODEX and DBiTplus images are aligned using tissue boundary detection and image transformation followed by the cell-to-spot mapping of CODEX and DBiTplus images and spot-level cell type deconvolution of the DBiTplus datasets using an RCTD-like approach. These spots are split to generate pure cell type subspots, and the two modalities are combined using Seurat WNN analysis.

### Development of Computational framework for DBiTplus

The premise of DBiTplus is to combine whole transcriptome sequencing (DBiT-seq) with high-plex protein imaging via CODEX on the same tissue slide. This approach facilitates precise cell-type identification at each spatial transcriptome spot, enabling image-guided decomposition to construct spatial transcriptome atlases at single-cell resolution. Practically, this would allow for the identification of the cell types of every single cell in the spatial transcriptome spot by leveraging the single-cell level protein information from the CODEX imaging of the same tissue slide. To achieve this, we developed a novel computational workflow that integrates the cell-by-protein matrix from CODEX and the spot-by-gene matrix from DBiT-seq to create a feature matrix with cell type labels and spatial locations from the two modalities (**Fig 1b)**. The workflow begins with whole cell segmentation of the CODEX dataset using DeepCell^36^ and quality control steps where the feature (protein marker) signals were summed, area-normalized, and scaled to a [0, 1] range based on the 5th and 95th percentiles, with values outside this range set to 0 or 1. Annotation of the entire CODEX dataset was achieved by integration with a reference scRNA-seq dataset, if available, such as the Mouse Organogenesis Cell Atlas Project^37^ (for mouse embryo samples), using MaxFuse which finds matched pairs between the scRNA-seq and CODEX datasets (**See Methods**). Utilizing tissue boundary detection and optimal image transformation, the CODEX and DBiT-seq images are registered within a common coordinate, which further enables accurate matching and identification of the same cells from the DBiTplus and CODEX datasets (**Extended Data Fig. 4e)**. To assign gene expression to specific cell types within DBiTplus spots, cell type proportions of the aligned CODEX cells are computed for each spot. Each spot is then divided into sub-spots based on pure cell types, allowing for cell-type-specific gene expression estimation. This approach enables the annotation of individual cells within each DBiTplus spot which is not possible from currently existing deconvolution tools (**Fig. 2g**). Approaches such as CARD^38^, RCTD^31^, SPOTlight^39^ and Cell2location^40^ typically rely on single-cell RNA-seq reference data or bulk cell-type expression profiles to estimate cell-type proportions across spatial coordinates as a pie chart. In contrast, our method directly utilizes information from the same-slice CODEX data for deconvolution, and then split spot-level spatial transcriptomic data into individual cell types, resulting in reduced bias and greater reliability. As demonstrated in **Extended Data Fig. 7**, our deconvolution method achieves superior cell type separation compared to TACCO when applied to DBiTplus data. By integrating DBiTplus and CODEX data using Seurat weighted-nearest-neighbor (WNN)^41^, we enhance cell type separation within the integrated embedding. Specifically, our deconvolution method, followed by Seurat WNN integration, successfully separates epithelial cells from other cell types in mouse embryos. In contrast, TACCO deconvolution fails to consistently identify the presence of epithelial cells in spatial spots when compared to CODEX data and loses these cells in the WNN embeddings.

**Fig. 2.**
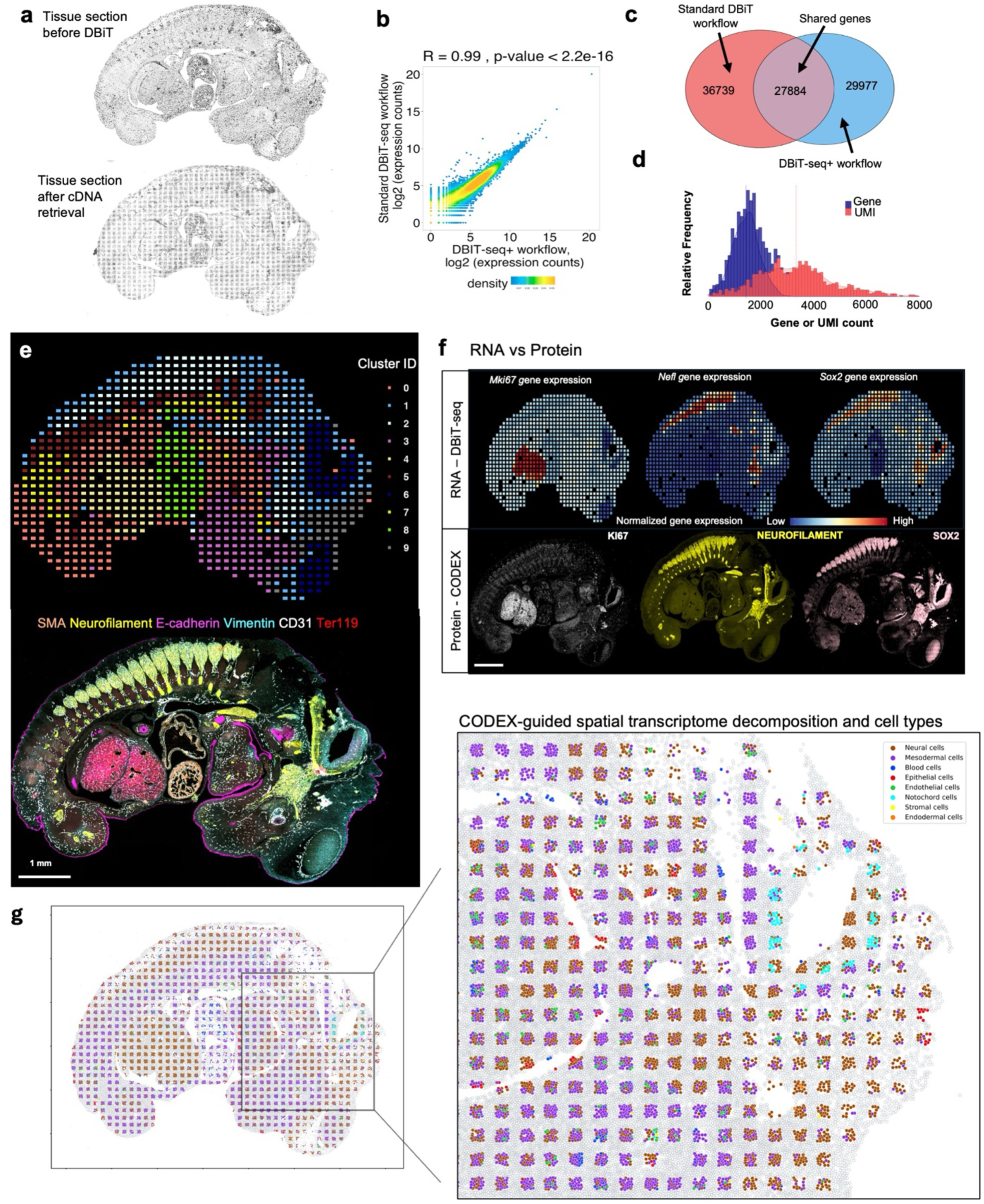
DBiTplus workflow, performance and spatial multi-omic analysis of FFPE mouse embryo. **a** Brightfield images E13 mouse embryo FFPE section used in the DBiTplus workflow. Left panel shows tissue section before the DBiTplus workflow and right panel shows the tissue section after cDNA retrieval with the tissue section intact and morphology preserved. **b** Correlation analysis between two samples (E13 mouse embryo FFPE section using standard DBiTplus workflow v E13 mouse embryo FFPE section using the DBiTplus+ workflow). Pearson correlation of 0.99 and p-value < 2.2e-16). **c** Venn diagram showing overlap of genes sequenced between the standard DBiTplus workflow and the DBiTplus workflow. Total of 27884 genes shared between the two datasets. **d** Distribution of detected gene/UMI counts per spatial spot. Dashed lines (blue and red) represent the average gene and UMI counts respectively. **e** Top: UMAP clustering identified 10 transcriptomic clusters of spots or spots from the E13 mouse embryo FFPE section. Bottom: Multiplexed immunofluorescence staining is performed on the same tissue section. **f** Comparison of the spatial gene expression from DBiTplus and protein expression from (CODEX). Similar localization observed in both modalities. Scale bar 1 mm. **g** CODEX-informed spot deconvolution of DBiTplus data. CODEX cells were annotated by label transfer from the MOCA dataset using MaxFuse matching. Annotated CODEX cells within each spot were used to inform composition of different cell types when performing deconvolution of DBiTplus spots using RCTD providing CODEX-informed average expression levels of different cell types in each spot.

### Workflow, development, and optimization in fresh frozen and FFPE samples

Preliminary experiments utilized OCT-frozen E13 mouse C57 embryo sagittal sections. Following the spatial barcoding process, the region of interest of the tissue was covered with a clean PDMS well gasket, and 50 μl of 0.1M NaOH was added atop the tissue and incubated at room temperature for 15 minutes, collected and neutralized with equimolar HCl. Imaging the tissue section revealed a disruption of the tissue morphology (**Extended Data Fig. 1b**). Using three serial sections of mouse spleen samples, one tissue section was subjected to cDNA retrieval using 0.1M NaOH for 5 minutes, the next section utilized 90% DMSO followed by an incubation of 30 minutes and the last section utilized 10U of RNaseH enzyme incubated at 37°C for 30 minutes after spatial barcoding. The samples were then sent for CODEX staining with a 25-marker mouse immune panel at Akoya Biosciences **(see Extended Data - Codex panel details**). Assessment of the quality of CODEX staining showed poor quality staining for both the tissue section with 90% DMSO and NaOH as the retrieval methods. However, the section utilizing RNaseH enzyme yielded 14 positively staining makers including CD45R (B cell lineage) and CD169 (macrophages) (**Extended Data Fig. 1c**). Thus, the enzymatic approach was selected for further optimization to enhance transcriptome and proteome data quality (**Extended Data Fig. 1d**).

To improve the method, a thermostable RNaseH enzyme, active above 37°C and optimal at over 65°C, was introduced. E13 mouse embryo sections were prepared, with one slide fixed in 1% formaldehyde (control) and the other in 0.5 formaldehyde% (test). Both slides underwent the same spatial barcoding process. The control slide was lysed with proteinase K following this while the test slide was incubated with 20U RNaseH at 55°C for 2 hours, and inactivated with 0.5M EDTA. Additional RNaseH was applied at 37°C overnight, followed by EDTA inactivation. The tissue was washed with 0.1X SSC buffer, and the collected cDNA was pooled, purified, template-switched, and PCR-amplified for sequencing. (**see Methods section**). Sequencing revealed 20,089 genes for the test slide versus 23,295 genes for the control, with the control showing about twice the mean gene count per spot, hinting at the cDNA release efficiency release via the enzymatic approach compared to lysis (**Extended Data Fig. 2a)**.

To further improve the cDNA release efficiency, 0.5% Triton X-100 was included in the RNaseH mix. The tissue integrity was maintained (**Extended Data Fig. 2b)**, and mean gene counts per spot were 1,029 (test) and 2,312 (control), respectively. A high correlation was observed between replicates and between test and control slides, (R=0.87) for both the first replicate and the second replicate (R=0.72). We also observed a strong correlation (R=0.84) between the test samples for both replicates (**Extended Data Fig. 2c**). Spatial clustering identified eight clusters, with clusters 2 and 3 being consistent with the embryonic diencephalon, mesencephalon and telencephalon, as evidenced by the expression of *Zbtb20^42^* and *Id4^43^* (embryonic neocortex, forebrain) from the heatmap of the top 5 differentially expressed genes from each cluster (**Extended Data Fig. 2d,e**). Spatial patterns of genes *Sox11* (regulation of progenitor and stem cell behavior including neurogenesis)^44^, *Col2a1* (cartilage primordium and vitreous humor of the eye)^45^, and *Sox2* (maintenance of the pluripotency of early embryonic cells)^46^, from clusters 2, 3 and 5 respectively, matched ISH data from the Allen Brain Atlas and that from spatially-enhanced resolution omics-sequencing from Stereo-seq^45^ (**Extended Data Fig. 2f).** Clusters 0 (control) and 2 (test) showed remarkably similar spatial distributions and significant overlap of 1109 genes, with gene ontology analysis revealing axogenesis as a top biological process associated with both clusters, underscoring DBiTplus’s effectiveness in generating high-quality spatial transcriptome data and recapitulating underlying tissue biology and structure.

We extended the workflow to FFPE tissues which is the ‘gold standard’ method for preserving clinical tissues with high-quality tissue morphology for clinical diagnosis^47^. Unlike fresh frozen tissues, FFPE tissues can be stored long-term at room temperature and retain tissue architecture and cellular morphology^48^. Thus, given the minimal quality of CODEX data obtained from prior fresh frozen tissues, we pivoted to FFPE tissues. Using the DBiTplus workflow, we profiled a human cerebellum and lymph node FFPE sections After preprocessing and cDNA retrieval, the tissue architecture remained intact (**Extended Data Fig. 3a,c**). The cDNA size ranged from 400-800 bp, typical for FFPE sections (**Extended Data Fig. 3d**). A 29-marker CODEX panel applied to the cerebellum (**see Extended Data - Codex panel details**) revealed distinct cerebellar layers, including the medulla, deeper granular cell layer, Purkinje cell layer, and molecular layer. Glial cells in the deep cerebellar medulla were identified by AQP4 staining, the single layer of large pear-shaped Purkinje cells, with their large cell bodies marked by calbindin could be visualized in the Purkinje cell layer (**Extended Data Fig. 3b**). CODEX staining of the lymph node showed the expected lymph node architecture, with CD20+ B cells and CD21+ follicular dendritic cells in defined follicles, and CD3ɛ+ T cells mainly in interfollicular areas alongside SMA+ smooth muscle cells (**Extended Data Fig. 3e-f**).

### Multimodal mapping of mouse embryo

The DBiTplus workflow, outlined in **Fig. 1a**, was performed on an E11 C57 paraffin-embedded mouse embryo, with the tissue section remaining intact after cDNA retrieval (**Fig. 2a**). A strong correlation (R=0.99) was observed between the DBiTplus workflow and the standard DBiT-seq workflow (on the adjacent section) with an overlap of 27,884 genes between the two datasets. On average, each spot captured approximately 1,200 genes and 3,300 UMIs (**Fig. 2b-d**). The intact section after DBiT was subsequently stained with a 26-marker CODEX panel (**see Extended Data - Codex panel details**). Unsupervised clustering identified 10 transcriptomic clusters, and the spatial clustering closely correlated with anatomic structures in the mouse embryo from the CODEX data (**Fig. 2e, Extended Data Fig. 4a-b, 5a**). From the heatmap, cluster 4 expressed *Hba.a1*, *Hba.a2*, *Hbb.bt*, *Afp* and *Serpina6* which are known to be associated with the embryonic liver^49^. Similarly, cluster 8 expresses *Myh6, Myl7, Myh7 and Tnnt2* which are known to be associated with the embryonic heart^50^ (**Extended Data Fig. 4c**).

We observed a similar spatial pattern of expression between mRNA transcripts and protein expression for *Mki67*, *Nefl* and *Sox2*, expressed in the (fetal liver and neurological system)^51^, (nervous system)^52^, and (neural tube^53^, retina^54^ and neocortex)^55^, respectively, as expected from E11 mouse embryos (**Fig. 2f**). Comparing the expression of three spatially distinct genes: *Sox2* (maintenance of the pluripotency of early embryonic cells)^46^ *Hoxa10* (limb muscles)^56^ and *Sox11* (regulation of progenitor and stem cell behavior including neurogenesis)^44^, from cluster 0, cluster 1 and a combination of clusters 1 and 2, respectively, to known in situ hybridization (ISH) spatial expression from Allen Brain Atlas and spatially-enhanced resolution omics-sequencing from Stereo-seq, we observed consistent spatial expression patterns, highlighting the capability of DBiTplus to accurately reflect underlying tissue biology and structure (**Extended Data Fig. 4d**), similar to results observed in OCT-frozen tissues. This underscores the technology’s versatility in effectively working with both FFPE and OCT-frozen tissues. A trained Support Vector Machine (SVM) model was applied to achieve the comprehensive cell type annotation of the whole CODEX dataset (**Extended Data Fig. 6a-d, 7a-d**). Following the integration of the CODEX and DBiT-seq datasets, we could annotate the individual cells within each specific DBiTplus spot with neural and notochord cells expressed in the brain region of the mouse embryo (**Fig. 2g**). As demonstrated in **Extended Data Fig. 8**, our deconvolution method achieves superior cell type separation compared to TACCO when applied to DBiTplus data. By integrating DBiTplus and CODEX data using Seurat weighted-nearest-neighbor (WNN)^41^, we enhance cell type separation within the integrated embedding. Specifically, our deconvolution method, followed by Seurat WNN integration, successfully separates epithelial cells from other cell types. In contrast, TACCO deconvolution fails to consistently identify the presence of epithelial cells in spatial spots when compared to CODEX data and loses these cells in the WNN embeddings.

### Multimodal mapping of normal human lymph nodes

We applied the DBiTplus workflow to reactive human lymph node samples. Adjacent 10 μm thick sections were used for these experiments with the 50 μm microfluidic device (**Fig. 3**) and 20 μm microfluidic device (**Extended Data Fig. 10**). After the cDNA retrieval process in the DBiTplus workflow, the tissue section was still intact and was then stained with a 35-plex CODEX panel. The flow cell was removed, and H&E staining was performed on the tissue section (**Fig. 3a, Extended Data Fig. 10a**). Unsupervised clustering revealed five distinct transcriptomic clusters, which closely corresponded with the spatial architecture and cell type distribution observed in the CODEX data. For example, from the heatmap of the top 5 differentially expressed genes for each cluster, cluster 1 expressed MS4A1 and CXCL13 which are associated with B cells, the corresponding CODEX image showed the same region highly positive for CD20. *Myh11* and *Cald1* expressed in cluster 3 are associated with smooth muscle actin and anatomically represent the medulla of this lymph node sample, which contains large blood vessels, sinuses, and medullary cords (**Fig. 3b, c, e**). Cluster 4 which is in close proximity to cluster 3, showed differential expression of MARCO (a macrophage receptor) expressed by macrophages lining the medullary cords^57^.

**Figure 3.**
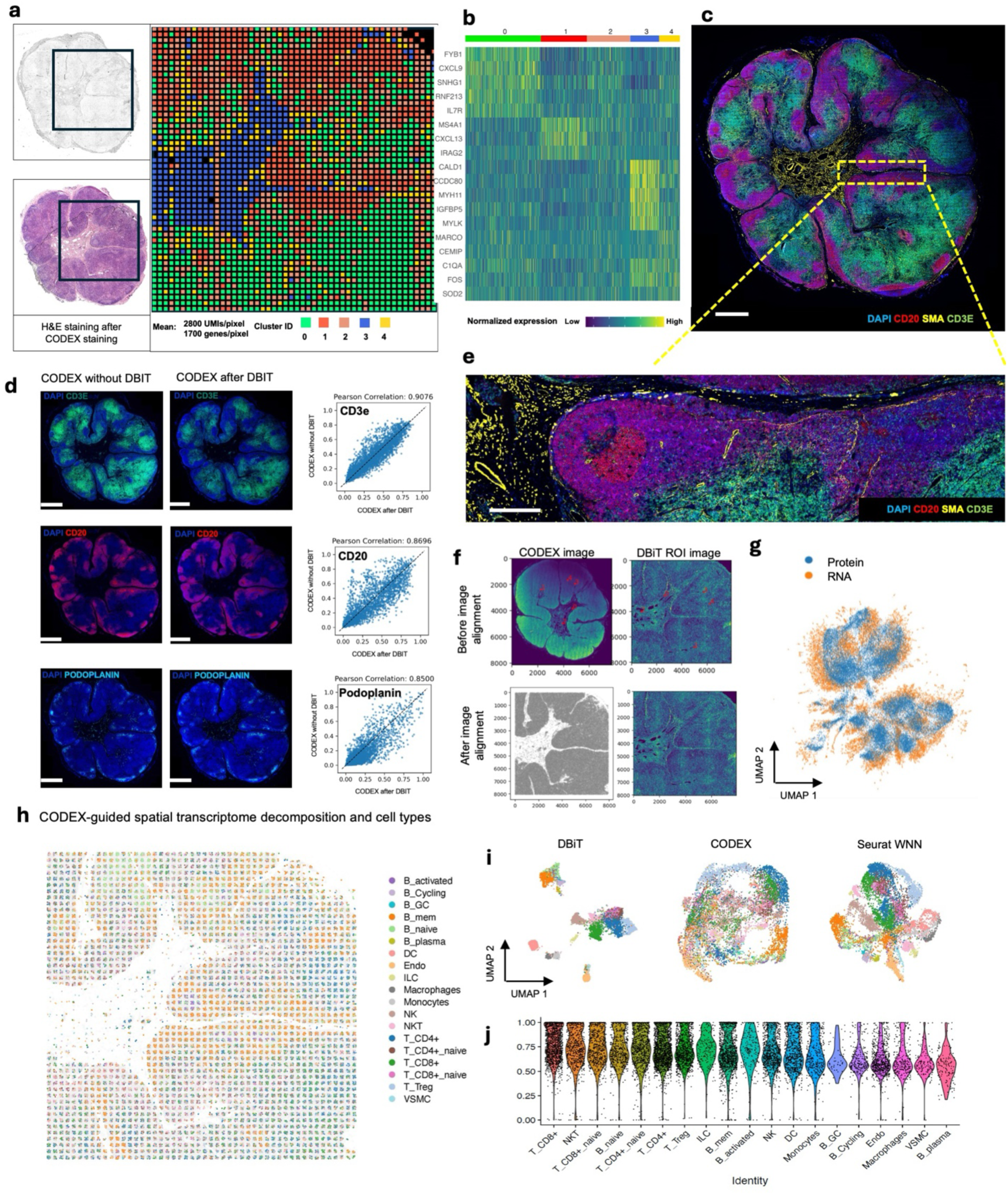
Spatial multi-omic profiling of FFPE human lymph node tissue section. **a** Spatial transcriptome distribution of FFPE human lymph node tissues barcoded using the microfluidic device with 50 μm resolution. Black box represents the DBiTplus ROI. UMAP clustering identified 5 distinct clusters whose spatial distribution closely matches with the histological organization of human lymph nodes. H&E staining is performed on same tissue after DBiTplus and CODEX. **b** Heatmap showing the top 5 differentially expressed genes for each spatial cluster. **c** CODEX staining (panel of 35 markers) is performed on the same tissue section following the DBiTplus workflow. Scale bar 1mm. **d** Correlation analysis between two adjacent sections stained via CODEX. Left: Tissue section stained via CODEX. Middle panel: Tissue section stained via CODEX after the DBiTplus workflow. Right: Scatter plot showing strong correlation in expression levels between the two samples. These plots confirm that high-quality CODEX data can be obtained from a tissue section following the DBiTplus workflow. Scale bar 1mm. **e** Zoomed in region from panel C from a representative area shows expression of CD20 (B cells) SMA (Smooth Muscles), and CD3ɛ (T cell marker). Scale bar 200 μm. **f** CODEX-DBiTplus image alignment using affine transformation and spatial landmarks in the two images. Top: Before image alignment. Bottom: After image alignment. **g** Integration of single-cell RNA seq reference data with CODEX data using the MaxFuse algorithm. Strong concordance observed between the two datasets. **h** Deconvolution of DBiTplus spots mapped by CODEX dataset. The identity of each cell within each DBiTplus spot can be directly inferred from CODEX staining on the same tissue section. **i** Seurat WNN integration of DBiTplus and CODEX datasets. **j** Weights of the CODEX modality of different cell types in Seurat WNN analysis. Strongest weights observed for T cell subtypes and lowest observed for B cell subtypes. These contributions are influenced by the composition of the CODEX marker panel.

To assess the quality of CODEX data obtained from a tissue section following the DBiTplus workflow, we imaged a lymph node tissue section using CODEX (located two sections away from the 50 μm sample) and performed correlation analysis for each marker. We observed a strong correlation for most major cell type markers, for example, CD3ɛ, CD20 and Podoplanin (Pearson correlations > 0.85) (**Fig. 3d**). Markers like CD45 and Beta-actin, which are broadly expressed across many cell types rather than being selectively expressed in specific ones, showed lower Pearson correlations (**Extended Data Fig. 9**). The DBiTplus and CODEX datasets were integrated using the workflow described earlier. The scRNA-seq dataset for human lymphoid tissues was obtained from the Cell2location repository^40^. UMAP for the joint embedding of the scRNA-seq dataset and CODEX dataset is shown in **Fig. 3g**. B cell subtypes (such as Germinal center B cells and naive B cells), T cell subtypes (such as CD8+ T cells and CD4+ T cells) can be observed in the follicular regions and interfollicular regions, respectively (**Extended Data Fig. 11 a, b**). The imprint of the DBiTplus channels on the tissue section can be seen in the Beta-actin staining from the CODEX image (**Extended Data Fig. 11c**) and SOX2 (**Extended Data Fig. 5b**) staining.

For both datasets obtained with the 25 μm and 50 μm pixel size experiments, the DBiTplus image was registered to the CODEX image to ensure that cellular-level data from the CODEX images were correctly aligned with the spatial transcriptomics data from the DBiTplus spots (**Fig. 3i, Extended Data Fig. 10d**). Assessment of the label prediction following label transfer showed high f1-scores for some cell types such as endothelial cells and monocytes with the lowest f1-score observed for germinal center B cells (**Extended Data Fig. 11d**). Each DBiTplus spot was divided into pure-cell-type sub-spots, which were then integrated with the CODEX cells within the spot by the Seurat WNN methodology (**Fig. 3h**). This approach identified each cell’s nearest neighbors based on a weighted combination of RNA and protein similarities. The cell-specific modality weights and multimodal neighbors were determined using the “FindMultiModalNeighbors” function. Higher ASW and ARI scores of 0.45 and 0.15, respectively, were observed with matched CODEX as deconvolution prior compared to deconvolution without CODEX data (**Extended Data Fig. 10e, f**). UMAP plots showed protein and RNAs for CD20 and CD21 were expressed by similar cell types, respectively in the joint embedding (**Extended Data Fig. 10g**). Violin plots showing the cell-specific modality weights from the WNN analysis indicated that the CODEX dataset provided an outsized importance for identifying most T cell subtypes compared to the DBiTplus data. However, the contribution of CODEX for the identification of B cell subtypes such as germinal center B cells and plasma B cells was lower, approximately, 0.55 (**Fig 3j**). This can be explained by the composition of the markers in the CODEX panel, which included more T cell markers than B cell markers (**see Extended Data - Codex panel details**). We assessed the quality of the H&E staining obtained after the DBiTplus workflow and one without any prior DBiT-seq or CODEX and observed comparable staining quality for both sets of images. While nuclear detail for the section after the DBiTplus workflow was lower, based on cell size and shape, as well as its location within the larger scaffold of the tissue, general cell type was still identifiable by trained pathologists. (**Extended Data Fig. 12**). As such, it is possible to utilize the histological image to infer single-cell spatial gene expression in the dead space regions between the microfluidic channels using newly developed tools like iStar^58^ and SPiRiT^59^.

### Multimodal mapping of human lymphoma transformation

Diffuse Large B Cell Lymphoma (DLBCL) is the most common lymphoma, representing about 30% of all such cases globally^60^. It is characterized by the rapid proliferation of large B-cells^61^. These lymphomas can be present in various sites, including lymph nodes and extranodal tissues, often leading to swift clinical deterioration if not promptly treated. DLBCL can arise “de novo” without any prior history of lymphoma diagnosis^62^. However, indolent lymphomas, such as follicular lymphoma (FL), marginal zone lymphoma (MZL), lymphoplasmacytic lymphoma (LPL), and chronic lymphocytic leukemia/small lymphocytic lymphoma (CLL), can potentially evolve into a more aggressive form such as DLBCL detected through histologic transformation, often with acquisition of MYC and BCL2 and/or BCL6 rearrangements^63^. Clinically, this transformation is often marked by a sudden and dramatic worsening of symptoms, including rapidly enlarging lymph nodes, systemic symptoms like fever, night sweats, and weight loss, and more aggressive disease behavior^64^. The prognosis following transformation is generally poor, with patients requiring more intensive therapeutic strategies. Around 60% of patients diagnosed with DLBCL can attain long-term remission and be cured following initial treatment with rituximab-based R-CHOP (rituximab, cyclophosphamide, doxorubicin, vincristine and prednisone) chemoimmunotherapy. However, autologous stem-cell transplantation (ASCT) is often considered necessary for some patients^61^. Using our DBiTplus workflow, we profiled a very interesting FFPE tissue sample, which is a DLBCL lymphoma arising from the transformation of MZL (**Fig. 4a**). Both BLBCL and MZL regions are observed in the same tissue section. Compared to the histology of healthy lymph nodes with well-defined follicles and interfollicular regions, DLBCL lymph nodes are characterized by the effacement of this normal tissue architecture by the infiltration of large, atypical B cells (**Fig. 4b**). We profiled a 2.5mm by 2.5mm region using our 25 μm microfluidic device (**Fig. 4c**) followed by CODEX on the same tissue section with a 44-marker panel (**Supplementary Table 2**). From the CODEX, we observed the diffuse arrangement of B cells marked by CD20 expression as well as patches of CD4 T helper cells (**Fig. 4d**). Integration of the DBiTplus transcriptomic data and the CODEX data allowed us to annotate and identify the cell type composition within the tumor microenvironment (**Fig. 4e, Extended Data Fig. 13a**). By comparing expression patterns on the UMAP of integrative embedding of cells using both protein and RNA information, we observed concordance in the expression of Vimentin, CD4 and CD20 in the joint embedding. While this has demonstrated the added to integrate DBiT spatial transcriptome and CODEX to identify cell types and zones, however, further analysis is warranted to further shed light on the tumor microenvironment in DLBCL, which is our ongoing work with samples from a cohort of patients.

**Figure 4.**
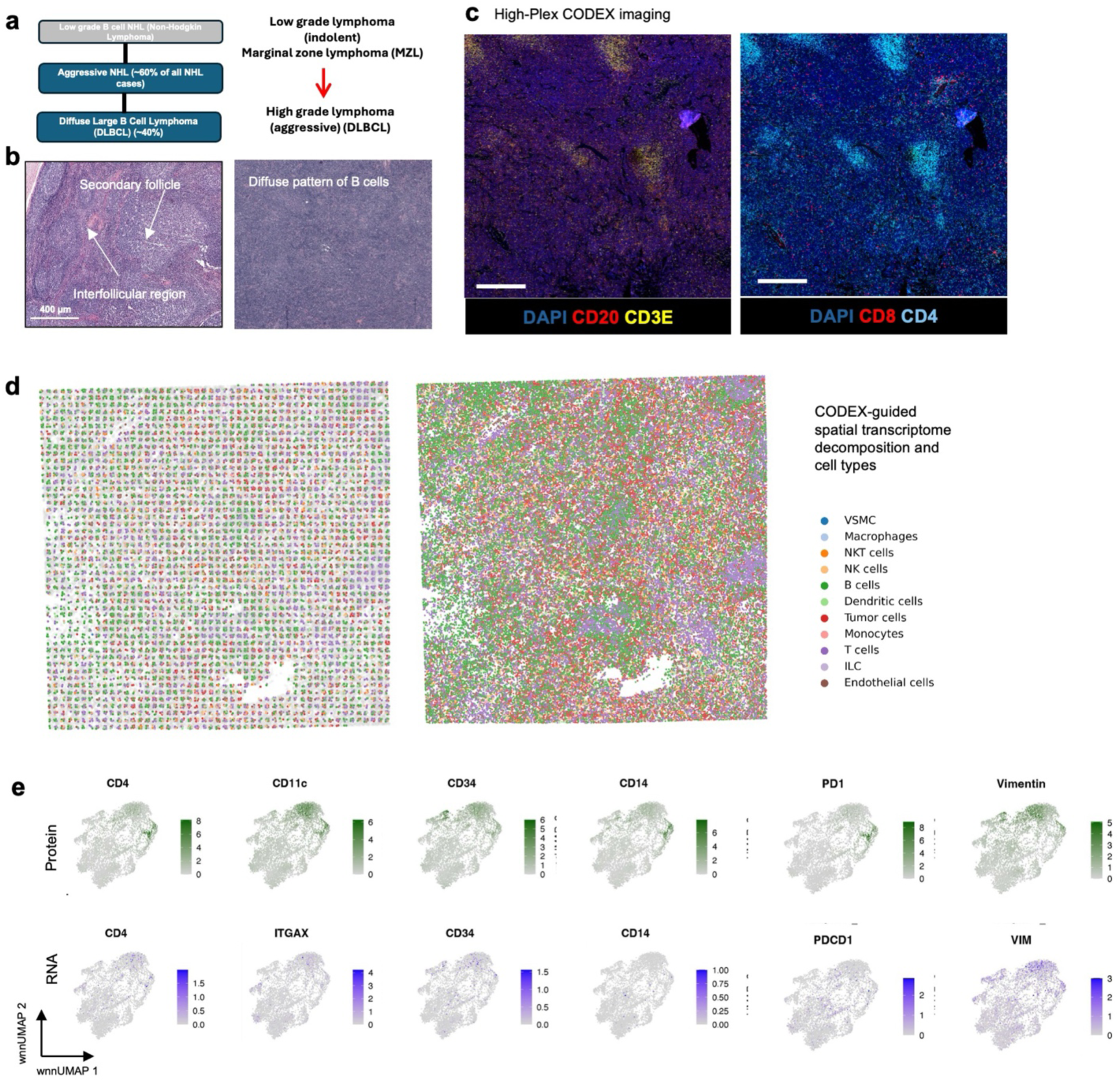
Spatial multi-omics profiling of FFPE human lymphoma tissue section. **a** Spatial analysis to unravel the molecular dynamics driving lymphoma transformation. Low-grade (indolent) marginal zone lymphoma transforms to high-grade (aggressive) DLBCL in patient with worse prognosis and outcomes. **b** H&E staining of normal FFPE lymph node and FFPE lymph node from patient with DLBCL. Left: Healthy lymph node showing distinct architectures of B cell and T cell zones. Right: DLBCL lymph node showing large diffuse distribution of B cells. Scale bar 400 um. **c** CODEX image of the same ROI profiled using the DBiTplus workflow. Left: CD20 (B cells) and CD3ɛ (T cells). B cells show diffuse pattern. Right: CD8 (Cytotoxic T cells) and CD4 (Helper T cells). Distinct patches of Helper T cells observed. Scale bar 200um. **d** CODEX-informed spatial deconvolution of cell types from the DBiTplus workflow. **e** Integration analysis of DBiTplus dataset (bottom panel) and CODEX dataset from FFPE DLBCL lymphoma (top panel).

## Discussion

Spatial multi-omic aims to integrate data from various omics layers—such as genomics, transcriptomics, proteomics, and metabolomics—while retaining the spatial context of biological interactions. This provides a comprehensive view of molecular processes, to help understand complex cellular structures and tissue microenvironments. Traditionally, spatial multi-omic approaches involve performing separate spatial omics assays on adjacent or serial tissue sections, followed by computational integration of these datasets. These approaches have been applied in studies of human myocardial infarction^24^ and the exploration of tumor microenvironments^26^. However, the inherent heterogeneity in cellular composition and tissue architecture between even closely adjacent sections makes it difficult to achieve precise alignment and integration of multimodal data from different tissue sections. Thus, new techniques for conducting multi-modal spatial multi-omics on the same tissue section are more desired. For example, MiP-seq was recently developed to detect multiplexed DNAs, RNAs, proteins, and biomolecules in brain tissue at subcellular resolution^28^ although the number of target molecules to map is rather limited or incompatible with unbiased molecular profiling at genome-scale.

In this study, we developed **D**eterministic **B**arcoding **i**n **T**issue **s**equencing **p**lus (abbreviated as DBiTplus) which combines unbiased transcriptome-wide spatially resolved sequencing with multiplexed immunofluorescence (CODEX) as well as novel computational pipelines for the profiling of the whole transcriptome and a panel of ∼50 protein markers on the same tissue section, compatible with both OCT-frozen and formalin-fixed paraffin-embedded (FFPE) tissues. Our experimental efforts have centered on optimizing the enzymatic cDNA retrieval process, which is a critical step. By using a Thermostable RNaseH enzyme, we were able to significantly improve the yield and quality of cDNA, while preserving tissue architecture. This chemistry workflow was pivotal as it directly impacted the ability to perform CODEX after DBiTplus, allowing for a comprehensive spatial analysis of gene and protein expression.

The optimized workflow demonstrated robust performance across different tissue types, including OCT-frozen mouse embryo sections and FFPE human lymph node and lymphoma sections. The comparative analysis of OCT-frozen tissues revealed that the enzymatic approach, particularly using the Thermostable RNaseH enzyme, resulted in higher gene capture efficiency and better tissue preservation compared to other tested methods such as the use of NaOH or DMSO. This was evidenced by the increased mean gene counts per spot and the strong correlation between replicates, indicating consistent and reproducible results. The spatial clustering of transcriptomic data further revealed distinct cellular structures, such as the embryonic brain regions, which were consistent with known anatomical features and gene expression patterns (**Fig. 2, Extended Data Fig. 2-5**). Transitioning to FFPE tissues, which are the gold standard in clinical diagnostics posed additional challenges. The formaldehyde-induced cross-linking in FFPE tissues often leads to RNA fragmentation and degradation, complicating spatial transcriptomic analyses. However, by adopting the patho-DBIT workflow to include RNA polyadenylation, we successfully profiled spatial gene expression in FFPE mouse embryos, human cerebellum and lymph node sections and lymphomas. Despite the inherent challenges associated with FFPE tissues, our workflow was able to maintain tissue integrity post-cDNA retrieval, as confirmed by both brightfield and DAPI imaging. The successful staining and imaging with CODEX panel, which identified key cellular markers and maintained the expected spatial architecture, underscore the robustness of our approach **(Extended Data Fig. 3**).

The integration of spatial transcriptomic data with high-resolution CODEX imaging on the same tissue slide represents a significant leap forward in spatial multi-omics. Computational innovation allows for data integration and construction of single-cell resolved spatial transcriptome atlases. First, by aligning the DBiTplus data with CODEX imaging through a refined registration process, we achieved precise co-localization of transcriptomic and proteomic data. Second, the co-localization enabled CODEX-informed DBiTplus spot deconvolution and the integration of transcriptomic information in deconvoluted spots with proteomic information in CODEX cells. This integration allowed for the accurate identification of cell types within the tissue, with cell-type-specific gene expression being assigned to individual cells in DBiTplus spots. Our approach, which involved splitting spots into pure-cell-type sub-spots and utilizing the Seurat weighted-nearest-neighbor (WNN) methodology, resulted in more reliable cell-type deconvolution, enriched feature dimensions, and provided a modality-integrated embedding for downstream analysis. Unlike previous methods based on blind deconvolution within each spot, using CODEX to guide the splitting of DBiT spatial transcriptome leads to truly single-cell level spatially resolved transcriptome atlases. The application of our DBiTplus workflow to human lymph node FFPE samples further demonstrated its versatility and applicability in clinical contexts. The observed spatial organization of lymph node architecture, with distinct clusters corresponding to known cellular regions, such as B cell-predominant follicles and T cell-rich zones, highlights the potential of this technology in studying complex tissues. The correlation between transcriptomic data and CODEX-based protein expression data also reiterates the accuracy and reliability of our multimodal integration approach.

However, DBiTplus shares common limitations with sequencing-based spatial transcriptomics approaches - low transcript capture depth can lead to a loss of information for genes with relatively low expression levels^65^. This issue is compounded by the fact that sequencing-based spatial transcriptomics methods generally rely on spatial barcoding and subsequent amplification processes, where any inefficiency in the capture or amplification steps disproportionately affects the detection of less abundant transcripts. This may be exacerbated by the potential lower transcript capture (release) rate for the enzymatic approach using RNaseH, used in DBiTplus compared to the tissue lysing approach in the standard workflow. This results in a loss of low-abundance transcripts, which can be particularly problematic in studies where these transcripts play critical roles, such as in the identification of rare cell types. In summary, while the DBiTplus method represents an advancement in spatial multi-omics by enabling high-resolution spatial mapping of the whole transcriptome and a panel of 50 proteins, its limitations in recovering low-abundance transcripts highlight the need for further technological improvements. These could include further optimization of the cDNA retrieval workflow (concentration of RNaseH enzyme, incubation times) to approximate the transcript release rate from the standard workflow, improved amplification methods, and strategies to increase overall capture efficiency to ensure a more comprehensive and unbiased representation of the transcriptome and proteome across different spatial contexts. Furthermore, the spot resolution of DBiTplus (demonstrated 20μm - 50 μm resolution) can be improved to approach single-cell resolution. However, one key advantage of DBiTplus is that by leveraging the protein data from the same slide, pure cell type composition of each spot can be generated. To demonstrate the versatility of our approach, future iterations can be paired with other spatial protein profiling technologies.

An alternative approach to our workflow that is being explored is to perform the multiplexed immunofluorescence (CODEX) workflow first, before the DBiT-seq workflow. This would preclude the need for the cDNA retrieval step using RNaseH. However, there are significant drawbacks to this approach since the repeated cyclic hybridization and washing steps in CODEX can strip weakly fixed mRNA transcripts within the tissue sections. The transcripts can also be susceptible to accelerated degradation and fragmentation from RNases.

In conclusion, the advancements in both experimental and computational innovation presented in this study significantly enhance the capabilities of DBiTplus for spatial multi-omics, making it a powerful tool for high-resolution spatial analysis across various tissue types, including those preserved by OCT and FFPE methods. The ability to integrate transcriptomic and proteomic data at a cellular level offers unprecedented insights into tissue architecture and cellular interactions, which are crucial for understanding complex biological systems and diseases. This work not only sets the stage for broader applications of spatially resolved multi-omics in research but also holds great promise for clinical diagnostics, particularly in the analysis of archived tissue samples.

## ACKNOWLEDGMENTS

We thank Yale West Campus cleanroom team for assistance with microfluidic wafer fabrications and the YPTS team for FFPE tissue sectioning and staining. Computational data analysis was conducted with Yale High Performance Computing clusters (HPC). We acknowledge the support received from the U.S. National Institutes of Health including grants U54CA274509, UH3CA257393, RF1MH128876, U54AG079759, U54AG076043, R01CA245313, RM1MH132648 (all to R.F.), and U01CA294514 (to R.F., M.X., and Z.M.). Z.M. is supported by NSF awards 2345215 and 2245575.

## Author Contributions

Conceptualization: R.F. and O.B. Methodology: A.E., H.Z., D.M., O.B., Z.Z. and R.F..; Experimental Investigation: A.E., H.Z., D.M., O.B., G.S., A.B. Data Analysis: A.E, Z.J., Z.B., M.Y., Y.L., and Z.M. Data Interpretation: M.L.X, R.F., O.B. and Z.M. Resources and valuable inputs: A.E, Z.Z., J.N, N.F., and N.Z. Original Draft: A.E., Z.Z., R.F. All authors reviewed, edited, and approved the manuscript.

## Materials and Methods

### Methods

#### Patient specimens

De-identified archived formalin-fixed paraffin-embedded (FFPE) human reactive lymph node and lymphoma tissue blocks were obtained from Yale Pathology Tissue Services (YPTS). The tissue retrieval and distribution for research was conducted with the approval of the Yale University Institutional Review Board and oversight by the Tissue Resource Oversight Committee. Written informed consent for participation in any cases where identification was collected alongside the specimen, was obtained from patients or their guardians, in accordance with the principles of the Declaration of Helsinki. Each sample was handled in strict compliance with HIPAA regulations, University Research Policies, Pathology Department diagnostic requirements, and Hospital by- laws.

#### Mouse tissue slides

Mouse C57 Embryo Sagittal Frozen Sections, E13 ([MF-104-13-C57](https://www.zyagen.com/product/mouse-c57-embryo-sagittal-frozen-sections-e13/)), Mouse C57 Spleen Frozen Sections ([MF-701-C57](https://www.zyagen.com/product/mouse-c57-spleen-frozen-sections/)) and Mouse C57 Embryo Sagittal Paraffin Sections, E11 ([MP-104-11-C57](https://www.zyagen.com/product/mouse-c57-whole-embryo-sagittal-paraffin-sections-e11/)) were purchased from Zyagen (San Diego, CA). Mouse Embryo Frozen Sagittal Sections, E13 and Mouse C57 Spleen Frozen Sections were made of freshly harvested tissues, snap frozen in OCT blocks and sectioned at a thickness of 7-10 μm and mounted on the center of poly-l-lysine-covered glass slides (63478-AS, Electron Microscopy Sciences). Mouse Whole Embryo Paraffin Sagittal Sections, E11 were made of freshly harvested tissues, fixed in 10% neutral buffered formalin, and processed for paraffin embedding. Paraffin blocks were also sectioned at a thickness of 7-10 μm and mounted on the center of poly-l-lysine-covered glass slides.

#### Human tissue section preparation

For human samples, after review of all slides by board-certified pathologist and selection of the optimal block, paraffin blocks were sectioned at a thickness of 7-10 μm and mounted on the center of Poly-L-Lysine coated 1 x 3" glass slides. Serial tissue sections were collected simultaneously for DBiT-seq and H&E staining. Human brain cerebellum paraffin sections ([HP-202](https://www.zyagen.com/product/human-brain-cerebellum-paraffin-sections-1604113886/)), were purchased from Zyagen and made of freshly harvested tissues, fixed in 10% neutral buffered formalin, and processed for paraffin embedding. Paraffin blocks of lymph nodes and lymphoma samples were also sectioned at a thickness of 7-10 μm and mounted on the center of poly-l-lysine-covered glass slides. The sectioning was carried out by YPTS. Paraffin sections were stored at −80°C until use.

#### Fabrication of microfluidic device

Details of the fabrication process for the PDMS wafers can be found in previous publications^66^. Customized high-resolution chrome photomasks were ordered from Front Range Photomasks and cleaned with acetone upon arrival to remove any contaminants. Master wafers were created by applying SU-8 negative photoresist (SU-2010 or SU-2025) to silicon wafers, adhering to the manufacturer’s instructions. Feature widths were set at 50 μm, 20 μm, or 10 μm, with corresponding heights of approximately 50 μm and 23 μm for the 50-μm-wide and 20-μm-wide devices. These wafers were then treated with chlorotrimethylsilane for 20 minutes to achieve high-fidelity hydrophobic surfaces. PDMS microfluidic chips were produced through replication molding, with the base and curing agents mixed in a 10:1 ratio and poured over the master wafers. Following a 30-minute degassing period in a vacuum, the PDMS was cured at 65-70°C for at least 2 hours. The cured PDMS slab was then cut out, and inlets and outlets were punched for further use.

#### DNA barcodes annealing

The DNA oligos used in this study were obtained from Integrated DNA Technologies (IDT, Coralville, IA), with the sequences provided in Supplementary Table 3 & 4. The barcodes (100 μM) and ligation linker (100 μM) were annealed at a 1:1 ratio in 2X annealing buffer (20 mM Tris-HCl pH 8.0, 100 mM NaCl, 2 mM EDTA). The mixes were placed in a thermal cycle and heated to 97°C to anneal and slowly cooled to room temperature at a rate of −0.1°C/s. The barcodes can be stored at −20°C for up to 6 months.

#### Tissue deparaffinization and decrosslinking

After the FFPE tissue section was taken from the -80°C freezer, it was allowed to equilibrate to room temperature. Then, the tissue slide was baked for 1 hour at 60°C to facilitate the removal of paraffin and increase adhesion of the tissue section to the slide. The tissue slide was then immersed into xylene twice for 5 minutes each to deparaffinize the tissue section. This was followed by rehydration steps in series of ethanol dilutions: two rounds of 100% ethanol, and one each of 90%, 70%, 50% and 30% ethanol. Finally, the tissue slide was immersed in distilled water for 5 minutes. Next, the tissue slide was immersed into pre-heated 1X antigen retrieval buffer and allowed to boil between 95-100°C for 10 minutes and then allowed to cool for approximately 30 minutes until it reached room temperature. The intact tissue slide was then imaged using the 10X objective on the EVOS M7000 Imaging System.

#### DBiTplus profiling of tissue

For OCT-embedded tissue sections stored in a −80 °C freezer, the tissue slides were allowed to equilibrate to room temperature. The section was then fixed with 4% formaldehyde for 20 minutes and washed three times with 0.5X DPBS-RI (1X DPBS diluted with nuclease-free water and 0.05 U/μl RNase Inhibitor. The tissue was permeabilized for 20 minutes at room temperature using 0.5% Triton X-100 in DPBS, followed by a wash with 0.5X DPBS-RI (1X DPBS diluted with nuclease-free water and 0.05 U/μl RNase Inhibitor) to stop the permeabilization. After air-drying, a PDMS reservoir was placed over the region of interest (ROI) on the tissue slide. In situ polyadenylation was performed with E. coli Poly(A) Polymerase. The samples were first equilibrated by adding 100 μl wash buffer (88 μl nuclease-free water, 10 μl 10X Poly(A) Reaction Buffer, 2 μl 40 U/μl RNase Inhibitor) and incubating at room temperature for 5 minutes. After removing the wash buffer, 60 μl of the Poly(A) enzymatic mix (38.4 μl nuclease-free water, 6 μl 10X Poly(A) Reaction Buffer, 6 μl 5U/μl Poly(A) Polymerase, 6 μl 10mM ATP, 2.4 μl 20 U/μl SUPERase•In RNase Inhibitor, 1.2 μl 40 U/μl RNase Inhibitor) was added to the reaction chamber and incubated in a humidified box at 37°C for 30 minutes. To remove excess reagents, the slide was dipped in 50 mL DPBS and shake-washed for 5 minutes. Next, 60 μl of the reverse transcription mix (20 μl 25 μM RT Primer, 16.3 μl 0.5X DPBS-RI, 12 μl 5X RT Buffer, 6 μl 200U/μl Maxima H Minus Reverse Transcriptase, 4.5 μl 10mM dNTPs, 0.8 μl 20 U/μl SUPERase•In RNase Inhibitor, 0.4 μl 40 U/μl RNase Inhibitor) was loaded into the PDMS reservoir and sealed with parafilm. The sample was incubated at room temperature for 30 minutes and then at 42°C for 90 minutes, followed by a wash with 50 mL DPBS. For the in-situ ligation of barcode A, the first PDMS device was carefully aligned over the tissue slide, positioning the 50 center channels over the ROI. The chip was imaged to record positions for downstream alignment and analysis. An acrylic clamp was used to secure the PDMS to the slide, preventing inter-channel leakage. The ligation mix (100 μl 1X NEBuffer 3.1, 61.3 μl nuclease-free water, 26 μl 10X T4 ligase buffer, 15 μl T4 DNA ligase, 5 μl 5% Triton X-100, 2 μl 40 U/μl RNase Inhibitor, and 0.7 μl 20 U/μl SUPERase•In RNase Inhibitor) was prepared. For the barcoding reaction, 5 μl of a ligation solution (comprising 4 μl of ligation mix and 1 μl of 25 μM DNA barcode A (A1-A50)) was added to each of the 50 inlets. The solution was drawn through the channels using a gently controlled vacuum. After incubating at 37°C for 30 minutes, the PDMS chip was removed, and the slide was washed with 50 mL of DPBS. Next, a second PDMS device with 50 perpendicular channels was attached to the air-dried slide over the region of interest (ROI). A bright-field image was taken, and barcode B was ligated similarly. The tissue section was then washed with nuclease-free water to remove any residual salts, and a final brightfield image was taken to mark the boundaries of the microfluidic channels on the tissue ROI.

#### cDNA extraction

For cDNA extraction, the region of interest of the tissue was covered with a clean PDMS well gasket, and 100 μl cDNA extraction solution (10 μl 5% Triton X-100, 74 μl nuclease-free water, 10μl 1X RNase H Reaction Buffer, and 6μl Thermostable RNaseH (M0523S, New England Biolabs)) was loaded into it. The reservoir was then clamped tightly with the slide to avoid any leakage and was sealed with parafilm. The clamped tissue slide was incubated in a humidified box at 55 °C for 3 hours. Following this, the cDNA extraction solution was collected and 1 μl of 0.5M EDTA was added to inactivate the RNaseH enzyme. The intact tissue slide was washed with 100μl of nuclease-free water which was then collected. Following this, 100 μl of cDNA extraction solution was added to the tissue slide as described as before and the clamped slide was incubated in a humidified box at 37°C overnight. The cDNA extraction solution was collected and 1μl of 0.5M EDTA was added to inactivate the RNaseH enzyme. The intact tissue slide was washed with 100μl of nuclease-free water which was also collected. The collected cDNA extraction solution was then pooled and kept in a -80°C freezer until use. For control slides, the standard lysing process described in previous publications from the lab was followed^66^.

#### DAPI staining

The tissue clamps were removed, and the intact tissue washed in nuclease-free water and then with 1X PBS. Following this, the tissue slide was incubated with 500μl of DAPI solution (two drops of NucBlue Fixed Cell ReadyProbes Reagent in 500 μl of 1X PBS) and incubated at room temperature for 5 minutes. The tissue slide was then imaged in the DAPI channel using the 20X objective on the EVOS M7000 Imaging System. This image would be used downstream to co-register the DBiTplus and CODEX images. The tissue slide was then washed with 1X PBS three times and stored at -80°C freezer until the CODEX imaging step.

#### cDNA purification, template switch, and PCR amplification

To construct the sequencing library, the pooled cDNA extraction solution was first purified using the Zymo DNA Clean & Concentrator-5 kit, following the recommended 5:1 ratio and eluted into 100 μl of nuclease-free water. Biotinylated cDNAs were captured using streptavidin beads (Dynabeads MyOne Streptavidin C1, Invitrogen). Prior to use, the beads were washed three times with 1× B&W buffer containing 0.05% Tween 20 and resuspended in 100 µl of 2× B&W buffer (10 mM Tris-HCl pH 7.5, 1 mM EDTA, 2 M NaCl). The beads were then mixed with the purified cDNA in a 1:1 volume ratio and incubated with gentle rotation at room temperature for 60 minutes. The beads were subsequently washed twice with 1X B&W buffer and once with 1X Tris buffer containing 0.1% Tween 20. Streptavidin beads bound with cDNA molecules were resuspended in 200 μl of TSO Mix (75 μl nuclease-free water, 40 μl 5X RT buffer, 40 μl 20% Ficoll PM-400, 20 μl 10mM dNTPs, 10 μl 200U/μl Maxima H Minus Reverse Transcriptase, 5 μl 40 U/μl RNase Inhibitor, 10 μl 100 μM TSO Primer). The template switch reaction was performed at room temperature for 30 minutes and then at 42°C for 90 minutes with gentle rotation. Afterwards, the beads underwent a single wash with 10 mM Tris-HCl pH 7.5 containing 0.1% Tween-20 and another wash with nuclease-free water. Second strand synthesis was then performed as follows: the beads were washed twice with TE-TW buffer (10 mM Tris pH 8, 1mM EDTA, 0.01% Tween 20) and resuspended in freshly prepared 200 μl 0.1M NaOH for 5 minutes with gentle rotation. The beads were washed once with 500 μl of TE-TW, and once with 500 μl 1xTE buffer (10 mM Tris pH 8, 1mM EDTA). The beads were then resuspended in 200 μl 2nd strand synthesis reaction solution (40 μl Maxima 5X RT Buffer, 20 μl 10 mM dNTPs, 2 μl 1 mM dN-SMRT oligo, 5 μl Klenow Enzyme 133 μl H2O) and rotated end-over-end at 37C for 1 hour. The beads were washed with nuclease-free water and were resuspended in 200 μl of PCR Mix (100 μl 2X KAPA HiFi HotStart ReadyMix, 84 μl nuclease-free water, 8 μl 10 μM PCR Primer 1, 8 μl 10 μM PCR Primer 2). This suspension was then distributed into PCR strip tubes. An initial amplification was performed with the following PCR program: 95°C for 3 minutes, five cycles at 98°C for 20 seconds, 63°C for 45 seconds, 72°C for 3 minutes, followed by an extension at 72°C for 3 minutes and a hold at 4°C. The PCR product was purified using SPRIselect beads at a 0.8X ratio, according to the manufacturer’s standard protocol. The resulting cDNA amplicon was then analyzed using the TapeStation system with D5000 DNA ScreenTape and reagents. The purified PCR product can be stored at -20°C until the next steps if necessary.

#### rRNA removal, library preparation, and sequencing

The SEQuoia RiboDepletion Kit was used to remove rRNA and mitochondrial rRNA from the amplified cDNA, following the manufacturer’s instructions. 20 ng of cDNA was used as the input amount, and three rounds of depletion were performed. We did observe that two rounds of depletion could suffice. Next, following the PCR steps described in the previous step, 10 PCR cycles was sufficient to directly ligate sequencing primers, using a 100 μl system consisting of 50 μl 2X KAPA HiFi HotStart ReadyMix, ∼42 μl solution from the rRNA removal step, 4 μl 10 μM Modified N501 Primer, and 4 μl 10 μM P7 Primer. The library product was purified using SPRIselect beads at a 0.8X ratio, according to the manufacturer’s standard protocol, and sent out to Novogene Corporation to be sequenced on an Illumina NovaSeq 6000 Sequencing System with paired-end 150bp read length.

#### CODEX spatial phenotyping using PhenoCycler-Fusion

A modified version of the CODEX PhenoCycler-Fusion protocol ([Link to protocol](https://www.akoyabio.com/wp-content/uploads/2021/01/CODEX-User-Manual.pdf)) was adopted for tissue sections used in the DBiTplus workflow. Since the tissue had already been deparaffinized and rehydrated during the DBiTplus workflow, the CODEX process began with a gentle antigen retrieval step using 1X AR9 buffer for 5-10 minutes. The tissue was then allowed to cool to room temperature and was rinsed twice with nuclease-free water and hydration buffer, followed by staining buffer as the antibody cocktail was prepared. The tissue slide was incubated with the antibody cocktail at room temperature for 3 hours in a humidity chamber. After incubation, the tissue underwent a series of steps including post-fixation, ice-cold methanol incubation, and a final fixation step. Attached to the flow cell, the tissue section was incubated in 1X PhenoCycler buffer with additive for at least 10 minutes to improve adhesion. The CODEX cycles were then set up, the reporter plate was prepared and loaded, and the imaging process began. A final qptiff file was generated, at the end which could be viewed using QuPath V0.5.1^67^. For further details on the PhenoCycler antibody panels, experimental cycle design, and reporter plate volumes, **see Extended Data - Codex panel details**.

#### Flow cell removal and H&E staining

Following the CODEX imaging workflow, the flow cell can be removed, and histological H&E staining performed on the same tissue section. To remove the flow cell, the tissue slide with the flow cell is immersed in xylene or HistoClear for a minimum of 20 minutes to weaken the adhesive. A razor was then used to carefully detach the flow cell from the tissue slide. The tissue slide was then rinsed thoroughly with deionized water and then with 1X PhenoCycler Buffer without additive three times for 10 minutes each. Histological H&E staining on the FFPE sections were conducted by YPTS.

#### Sequence alignment and generation of gene expression matrix

Read 2 from the FASTQ file was processed to extract unique molecular identifiers (UMIs) and spatial Barcodes A and B. Read 1, containing cDNA sequences, was aligned to the mouse GRCm39 or human GRCh38 reference genome using STAR V2.7.8a^68^. Spatial barcode sequences were demultiplexed with ST_Pipeline V1.8.1^69^, using the predefined coordinates of the microfluidic channels, and ENSEMBL IDs were converted to gene names. This generated a gene-by-spot expression matrix for downstream analysis. Entries in the matrix that corresponded to spot positions without tissue were excluded.

#### Gene data normalization and unsupervised clustering analysis

Spatial gene expression analysis was conducted using the Seurat V4.3.0 pipeline^41^. Initially, gene expression within each spot was normalized and variance-stabilized using the SCTransform method, specifically designed for single-cell RNA sequencing (scRNA-seq) datasets. Linear dimensional reduction was then performed with the "RunPCA" function, and the optimal number of principal components for further analysis was determined using a heuristic approach, visualized by an ‘Elbow plot’ that ranks PCA components by their variance percentages. Subsequently, the "FindNeighbors" function embedded spots into a K-nearest neighbor graph structure based on Euclidean distance in PCA space, and the "FindClusters" function applied a modularity optimization technique to cluster the spots. The "RunUMAP" function was used to visually explore spatial heterogeneities through the Uniform Manifold Approximation and Projection (UMAP) algorithm. Finally, differentially expressed genes (DEGs) defining each cluster were identified using the "FindMarkers" function for pairwise comparisons between spot groups.

#### Single-cell RNA-seq reference data

All single-cell RNA sequencing (scRNA-seq) data utilized in this study are publicly available. The scRNA-seq dataset for the mouse embryo was obtained from the Mouse Organogenesis Cell Atlas Project, which can be accessed at [Link to MOCA dataset](https://oncoscape.v3.sttrcancer.org/atlas.gs.washington.edu.mouse.rna/landing). For this study, cells with a "development_stage" of 10.5 to 11.5 days post-coitum were selected to correspond with CODEX measurements, using the MaxFuse method as described below. Doublet removal and cell annotations were conducted by the original authors of the dataset. The scRNA-seq dataset for human lymph nodes was retrieved from the Cell2location repository, and is available for download at Link to reference dataset. Cell type annotations were provided by the original study.

#### Preprocessing of CODEX data

Whole-cell segmentation was performed using the Mesmer method^36^ employing pretrained weights. The DAPI channel served as the nuclear marker, while the CD45 channel was utilized as the membrane marker. For cell mask prediction, the default training resolution of 0.5 μm per spot was adopted. Cells within the [0.05, 0.95] cell-size quantile range and possessing DAPI signal intensities exceeding the 0.1 quantile threshold were retained for analysis. CODEX features were extracted by summing the signal for each feature per cell. For each CODEX feature, the 0.05 and 0.95 quantiles were calculated, and each single-cell level CODEX feature was subsequently scaled to a [0, 1] range, with the 0.05 quantile mapped to 0 and the 0.95 quantile mapped to 1. Values exceeding this range were clipped to 0 or 1, as appropriate.

#### Integration of CODEX with scRNA-seq dataset

The integration of scRNA-seq data and CODEX data was accomplished utilizing the MaxFuse algorithm^30^. Prior to integration, standard preprocessing protocols were applied to all scRNA-seq data using Scanpy. This preprocessing included count normalization, log1p transformation, and the identification of highly variable genes, resulting in the selection of 5000 genes exhibiting the highest variability. Linked features between the scRNA-seq and CODEX datasets were identified based on corresponding protein and gene names. From these linked features, those with a standard deviation greater than 0.01 were selected to enhance integration performance. During the pivot matching process, the number of principal components used to construct the nearest neighbor graph was determined by examining the elbow of the singular value decomposition (SVD) plot. A smoothing weight of 0.3 was applied, as recommended by the MaxFuse method, to account for the weak linkage between the two modalities. Following integration, low-quality pivots were removed to ensure the reliability of the cross-modal pivot pairs. Approximately 10% of cells from the CODEX datasets, representing high-confidence matches, were selected to construct these pivot pairs. For each pivot pair, cell type labels from the scRNA-seq data were transferred to the matched CODEX cells. To extend cell type annotation to the entire CODEX dataset, a Support Vector Machine (SVM) model was trained on the pivot CODEX cells to predict cell type labels based on protein expression measurements. Once trained, the model was applied to the remaining non-pivot cells within the CODEX dataset, thereby achieving comprehensive cell type annotation across the dataset.

#### CODEX-DBIT alignment

To align the high-resolution CODEX image with the low-resolution DBiTplus spot image of mouse embryo dataset, tissue boundaries were initially identified in both images. The input image was first smoothed using a Gaussian filter (implemented via the “filters.gaussian” function from the “skimage” package) to reduce noise and enhance relevant structures. Subsequently, Otsu’s thresholding method (employed through the “filters.threshold_otsu” function from the “skimage” package) was applied to the smoothed image to compute an optimal threshold value. This threshold was then used to generate a binary image, effectively separating the foreground (potential tissue) from the background. To ensure the detected regions were solid, any holes within the binary regions were filled using a binary hole-filling algorithm (utilizing the “binary_fill_holes” function from the “scipy” package). The final output was a binary image with no holes enabling identification of the tissue’s outer boundary. Following the identification of tissue boundaries, an optimal similarity transformation was determined and applied to the CODEX image. This transformation was computed using the “transform” module from the “skimage” package, aiming to minimize the squared error loss between the transformed CODEX image and the DBiTplus image. Given the cell positions in the CODEX image and spot positions in the DBiTplus image, the learned image transformation can be utilized to register the cells. This registration ensures that cellular-level data from the CODEX images are correctly aligned with the spatial transcriptomics data from the DBiTplus spots.

#### DBiTplus spot cell type deconvolution and splitting

The deconvolution of DBiTplus spots is achieved by computing the cell type proportions of CODEX cells aligned to each spot. Let *β_i,k_* denote the proportion of the contribution of cell type *k* to spot *i*, *x_i,j_* denote the total expression of gene *j* in DBiTplus spot *i*, and *μ_k,j_* denote the average expression of gene *j* of cell type *k* in the scRNA-seq data. To attribute gene expression to specific cell types within each DBiTplus spot, we split each spot into pure-cell-type sub-spots. This is done by computing the expected cell-type-specific gene expression at each DBiTplus spot. The calculation uses the following formula:

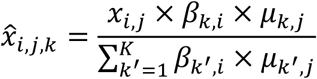

#### Seurat WNN analysis

CODEX cells were aggregated into pure-cell-type sub-spots, and DBiTplus spots were similarly divided into pure-cell-type sub-spots. Subsequent analyses were performed using the Seurat weighted nearest neighbor (WNN) methodology, with measurements from the pure-cell-type sub-spots serving as input data. The RNA data underwent normalization using a scale factor derived from the median unique molecular identifier (UMI) count. Following normalization, variable features were identified, and principal component analysis (PCA) was conducted on the RNA assay. Subsequently, the default assay was switched to the protein modality, variable features were identified, and PCA was performed on the protein data under the reduction name ‘apca’. For each cell, the nearest neighbors within the dataset were determined based on a weighted combination of RNA and protein similarities. The cell-specific modality weights and multimodal neighbors were calculated using the “FindMultiModalNeighbors” function. The integrated results were then utilized for visualization and clustering.

#### Assessing quality of CODEX data after DBiT-seq

The standard CODEX data preprocessing and filtering pipeline was first applied to extract cell positions and cell-level features from two adjacent CODEX slices separately. Six marker points were then manually identified from the two stacked CODEX images. The optimal affine transformation, which minimized the distance between paired marker points, was estimated using the "transform" module from the Python package “skimage”. This transformation was subsequently applied to align cell positions from the two slices into a common coordinate system. To ensure statistical robustness, the spatial dimensions were discretized into bins with a pixel size of 100 (∼4000 common bins), and the mean values of selected features were computed by aggregating the cells within each bin. Common bins across both datasets were identified, and Pearson correlation coefficients were calculated to quantify the strength of linear relationships between corresponding features, and scatter plots were generated to visually represent the correlations.

## EXTENDED DATA FIGURES

**Extended Data Fig. 1.**
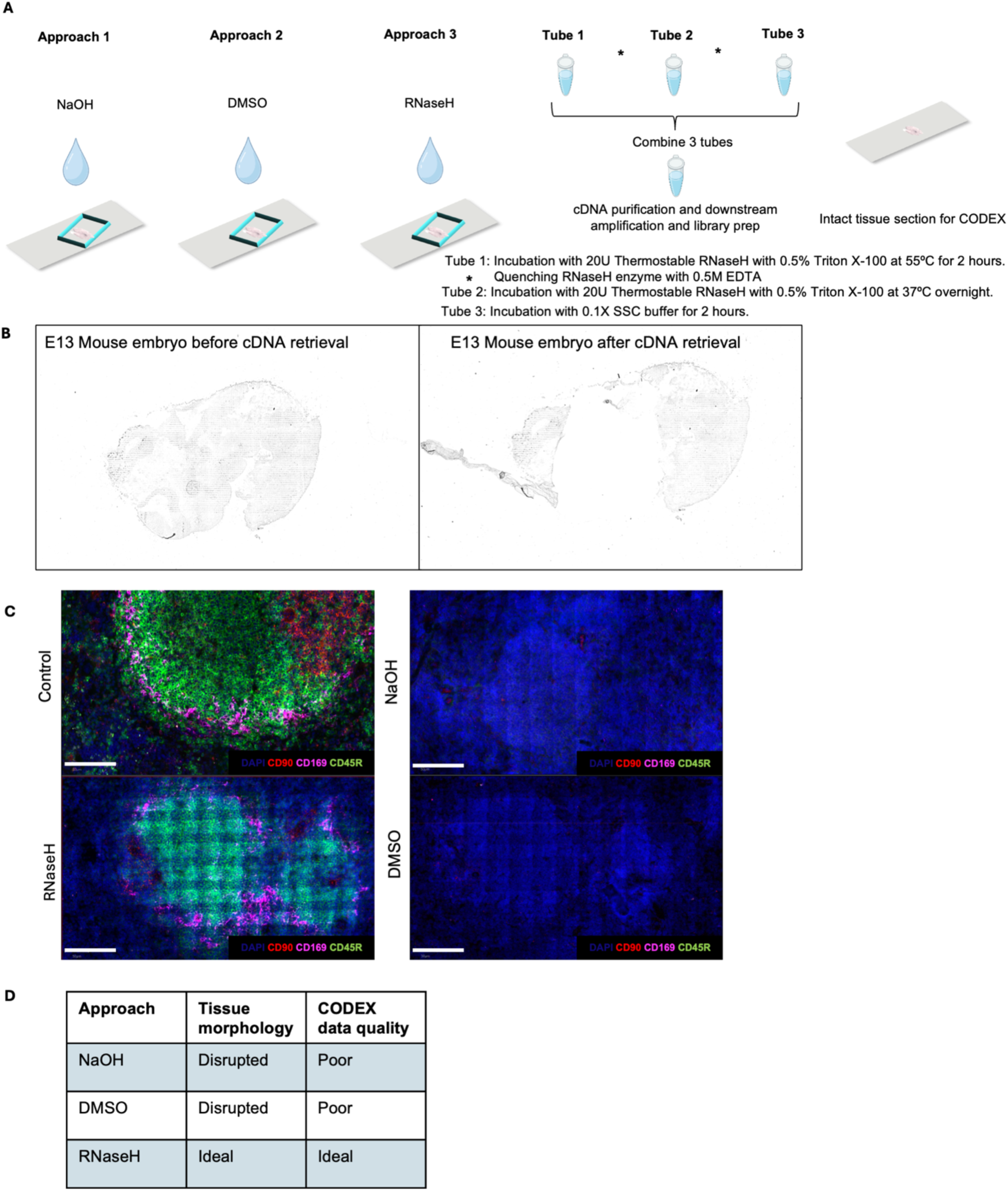
Exploration of different approaches for cDNA retrieval from FF and FFPE tissue sections. **a** Approaches for releasing cDNA from tissue section. Three approaches were tried – NaOH, DMSO (which can allow for the dissociation of cDNA followed by diffusion from the tissue section) and RNaseH enzyme (can selectively degrade mRNA in a DNA:RNA hybrid followed by diffusion from the tissue). Thermostable RNaseH allows for incubation at higher temperatures to facilitate diffusion. Three tubes are then pooled, cDNA purification and downstream amplification and library prep is done. The intact tissue section is then used for CODEX staining. **b** Brightfield images E11 mouse embryo FF section used in the DBiTplus workflow. Left panel shows barcoded tissue section after the DBiTplus workflow and right panel shows the barcoded tissue section after the cDNA retrieval. **c** Comparing the quality of CODEX imaging on FF mouse spleen after DBiTplus workflow. Mouse spleen sample with the retrieval method using RNaseH enzyme shows positive staining makers including CD45R (B cell lineage) and CD169 (macrophages) similar to control staining. Scale bar 50 μm. **d** QC analysis of parameters for selection of optimal approach for cDNA retrieval from the tissue section. RNaseH approach preserves tissue morphology and generates good quality CODEX data.

**Extended Data Fig. 2.**
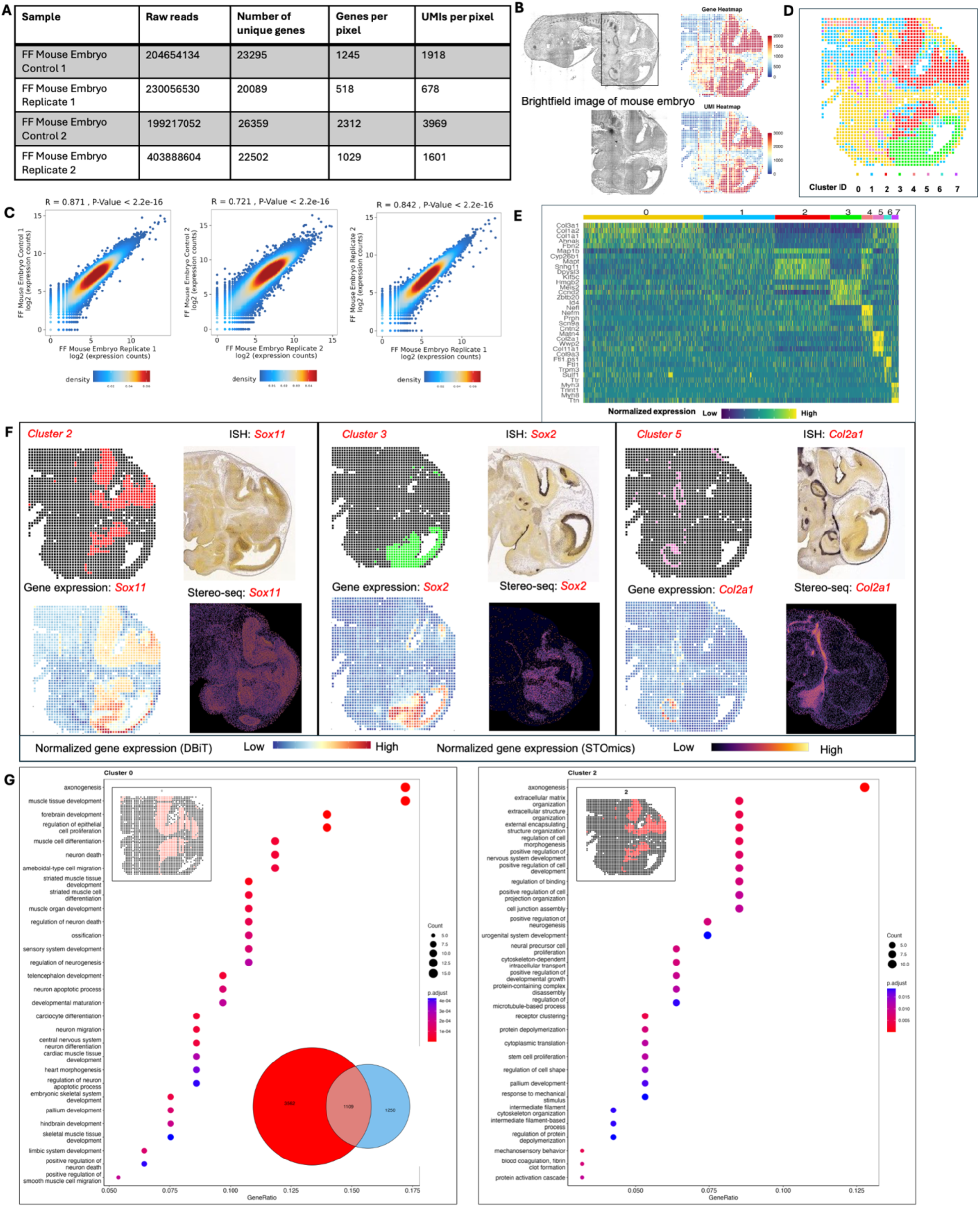
Spatial profiling of gene expression in FF mouse embryo. **a** Summary of raw reads, total number of unique genes, and genes and UMIs per 20 μm spot for FF mouse embryo experiments. **b** Brightfield image of E11 FF mouse embryo. Black box represents the DBiTplus workflow. Spatial mRNA and UMI count maps. An average of 1029 genes and 1601 UMIs detected per 20 μm spot. **c** Correlation analysis between replicates showing the reproducibility of the DBiTplus workflow with a Pearson correlation coefficient of 0.87 and 0.72 respectively. Pearson correlation between the two replicates is 0.84. **d** Spatial UMAP showing 8 distinct clusters of spots or spots that align with anatomical features of the E11 mouse embryo. **e** Heatmap showing the top 5 differentially expressed genes for each spatial cluster. **f** Spatial distribution of clusters 2, 3 and 5 alongside their principal defining genes. ISH staining image from the Allen Mouse Brain Atlas and gene expression from Stereo-seq data. **g** GO analysis showing the biological process associated with cluster 0 (E11 mouse embryo from the standard DBiTplus workflow) and cluster 2 (E11 mouse embryo from DBiTplus workflow) which are associated with the embryonic diencephalon. Venn diagram showing overlap of genes between cluster 0 and cluster 2.

**Extended Data Fig. 3.**
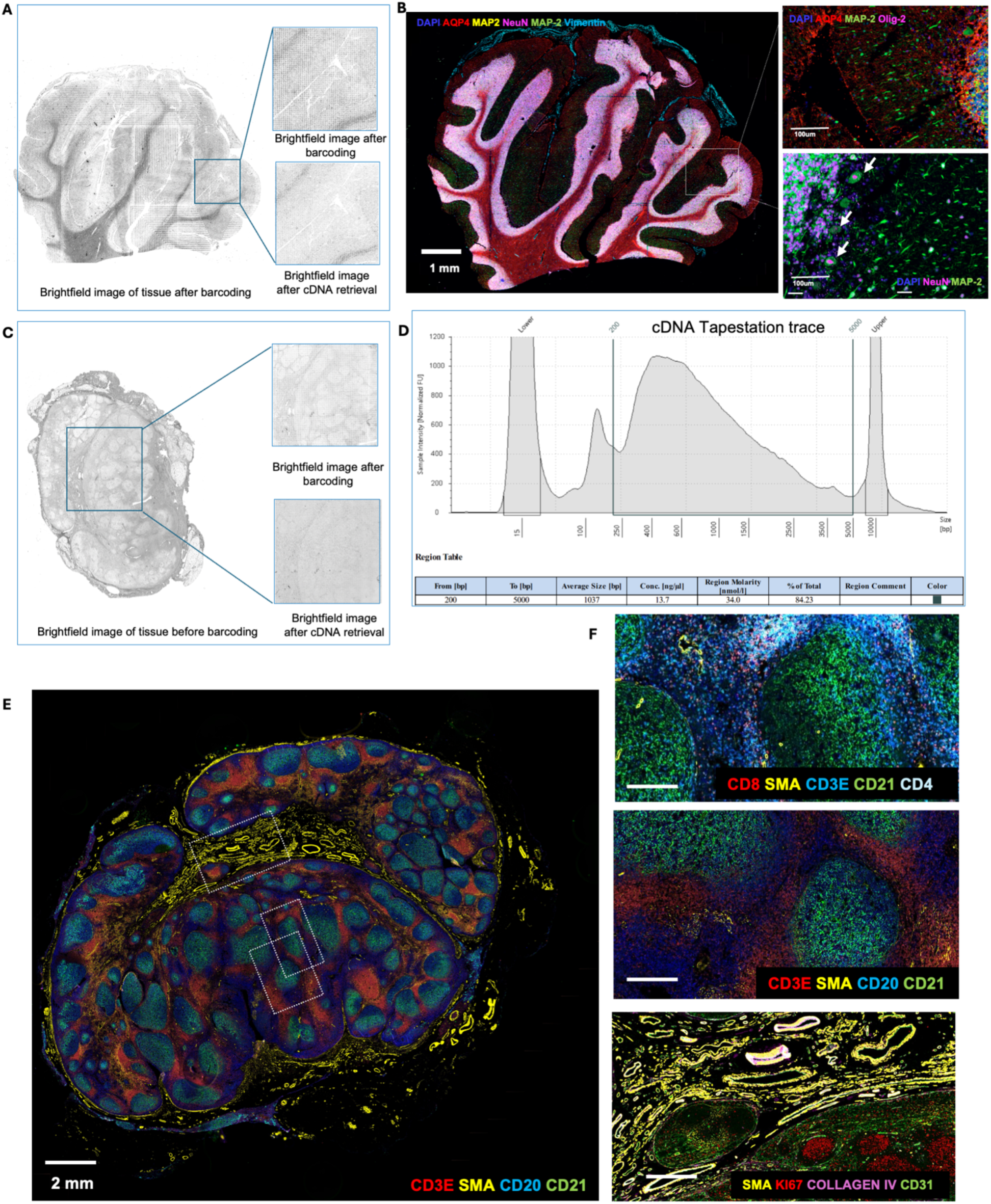
Optimization of CODEX staining on FFPE human cerebellum and lymph node following the DBiTplus workflow and cDNA retrieval using RNaseH. **a** Brightfield image of FFPE human cerebellum following barcoding in the DBiTplus workflow. Box represents the barcoded region. Brightfield image after cDNA retrieval shows the tissue morphology is preserved and is suitable for CODEX imaging. **b** The same tissue section from panel A is imaged with a 28-marker panel. Left: CODEX image showing AQP4 (Astrocytes), MAP2 (Synaptic neurons), NeuN (Neuronal nuclei and cell bodies) and Vimentin (Vasculature). Right top: Synaptic neurons marked by MAP2 in the molecular layer of the cerebellum. Right bottom: Large pear-shaped Purkinje cells, with their large cell bodies visualized in the Purkinje cell layer. **c** Brightfield image of FFPE of benign human lymph node. Box represents the barcoded region. Brightfield image after cDNA retrieval shows the tissue morphology is preserved and is suitable for CODEX imaging. **d** Size distribution from TapeStation traces of cDNA amplicon. **e** CODEX imaging on FFPE of human lymph node following the DBiTplus workflow. Three distinct regions are further analyzed for quality of CODEX staining. **f** Top: Region showing T cell subtypes: CD3ɛ (T cells), CD8 (Cytotoxic T cells), CD4 (Helper T cells) and SMA (Smooth muscles). Middle: CD3ɛ (T cells), SMA (Smooth muscles), CD20 (B cells) and CD21 (Follicular dendritic cells). Bottom: SMA (Smooth muscles), Ki67 (Proliferating germinal center B cells), Collagen IV (Blood vessels), CD31 (Vasculature).

**Extended Data Fig. 4.**
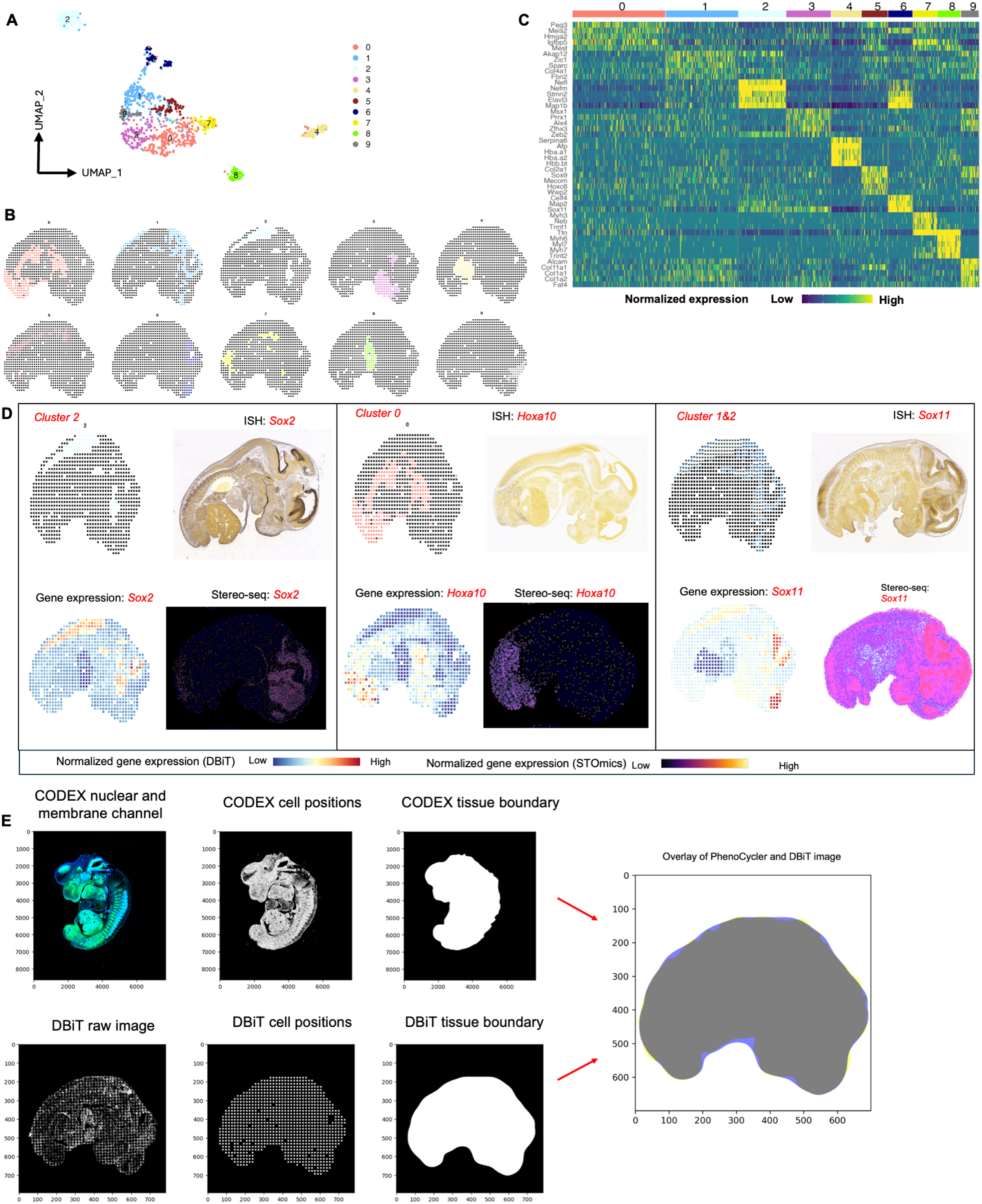
Workflow for the alignment of CODEX and DBiTplus images. **a** UMAP showing 10 distinct clusters of DBiTplus spots/spots. **B** Spatial distribution of the 10 clusters of DBiTplus spots. **c** Heatmap showing the top 5 differentially expressed genes for each spatial cluster. **d** Spatial distribution of clusters 0, 1, and 2 alongside their principal defining genes. ISH staining image from the Allen Mouse Brain Atlas and gene expression from Stereo-seq data. **e** CODEX-DBiTplus image alignment workflow. Top: Cell segmentation based on nuclear marker (DAPI) and two membrane markers, followed by determination of the coordinates of each cell in the CODEX dataset and tissue boundary detection. Bottom: Coordinates of the spots (spots) is extracted from the DBiTplus image, and the boundaries of the tissue section is detected. The CODEX image and the DBiTplus images are overlayed and registered in a common coordinate system.

**Extended Data Fig. 5.**
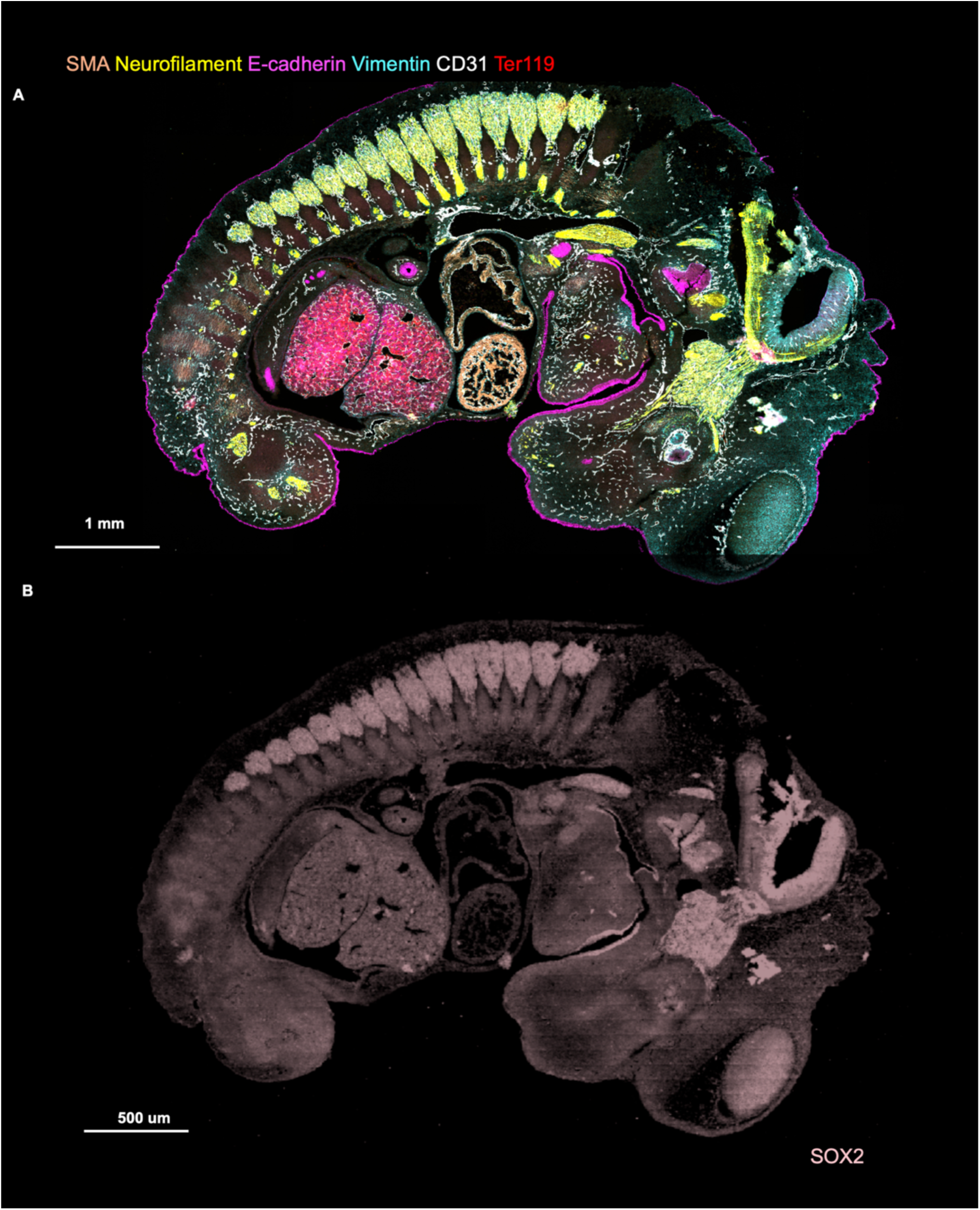
CODEX imaging of E13 FFPE mouse embryo. **a** Following the DBiTplus workflow, the intact tissue section is stained with a 21-marker panel. E-cadherin (Epithelia), CD31 (Vasculature), SMA (Smooth muscle), Ter119 (Erythroid cells), Vimentin (Mesenchyme and Endothelial cells) and Neurofilament (Nervous system). **b** Imprint of DBiTplus microfluidic channel on the tissue section observed in the SOX2 channel.

**Extended Data Fig. 6.**
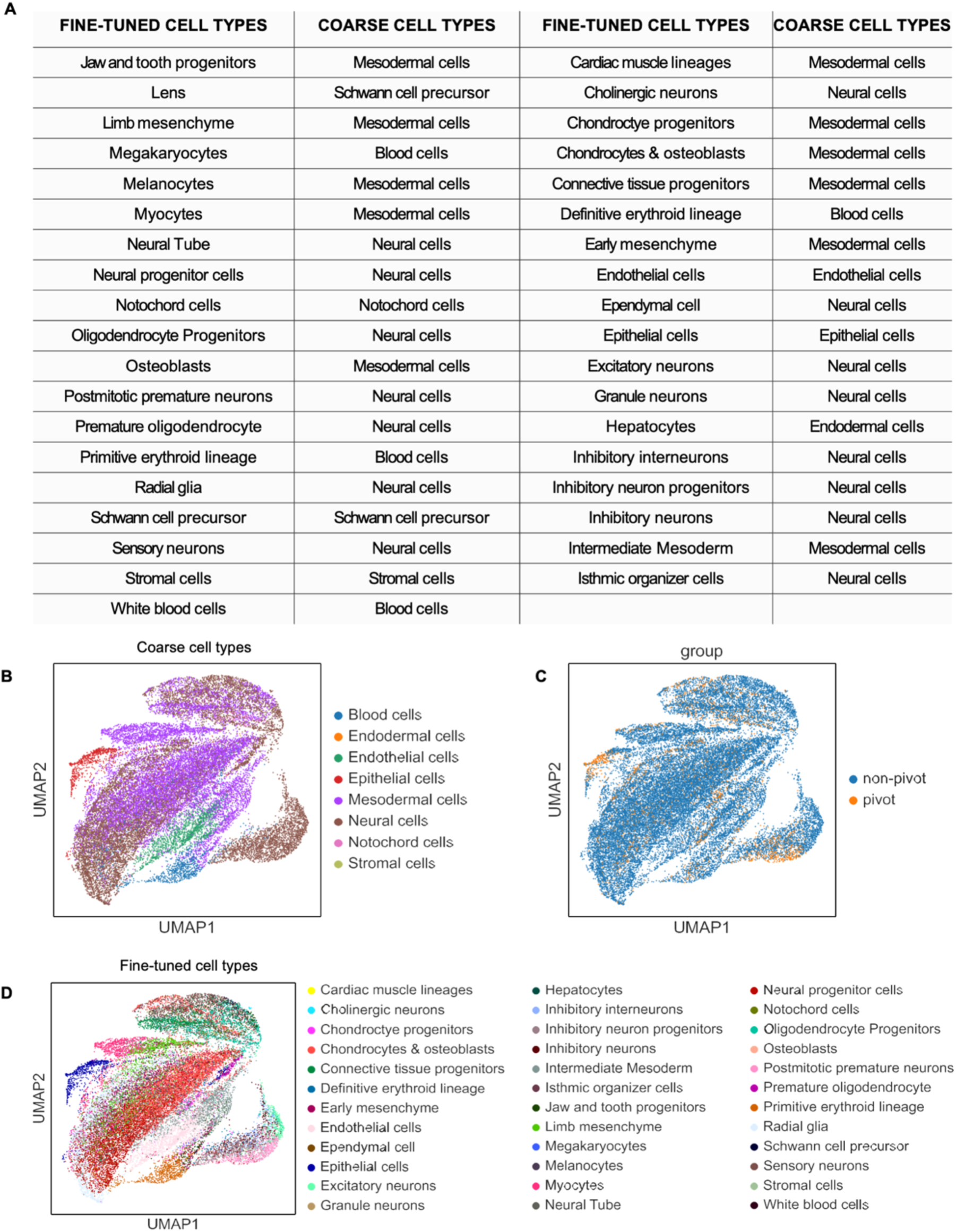
UMAP of integration of CODEX dataset and single cell RNA-seq reference dataset from E13 FFPE mouse embryo using MaxFuse. **a** Table showing the coarse and fine-tuned cell types from the single cell RNA-seq reference dataset. **b** UMAP showing major cell types from the integration of the single cell RNA-seq reference dataset and CODEX. **c** UMAP of the pivot and non-pivot cells in the MaxFuse integrated dataset. **d** UMAP of minor cell types in the integrated dataset.

**Extended Data Fig. 7.**
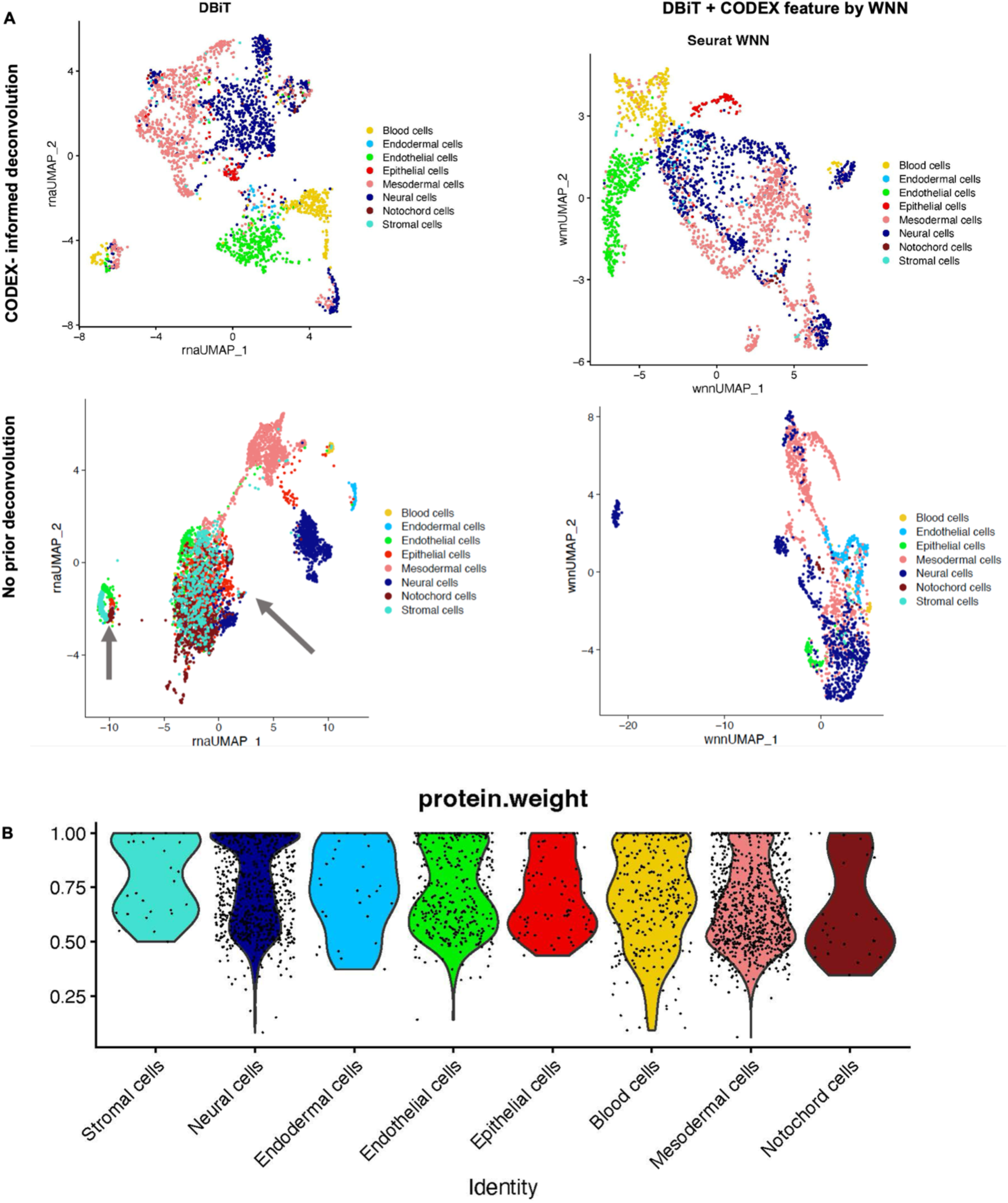
Integration of CODEX dataset and single cell RNA-seq reference dataset from E13 FFPE mouse embryo using MaxFuse. **a** Spatial plot of pivot cells and all cells from the matched single cell RNA-seq and CODEX datasets for major cell types. **b** Spatial plot of pivot cells and all cells from the matched single cell RNA-seq and CODEX datasets for minor cell types. **c** Spatial plot of pivot cells and all cells from the matched single cell RNA-seq and CODEX datasets for minor cell types belonging to neural cells. **d** Spatial plot of pivot cells and all cells from the matched single cell RNA-seq and CODEX datasets for minor cell types belonging to mesodermal cells.

**Extended Data Fig. 8.**
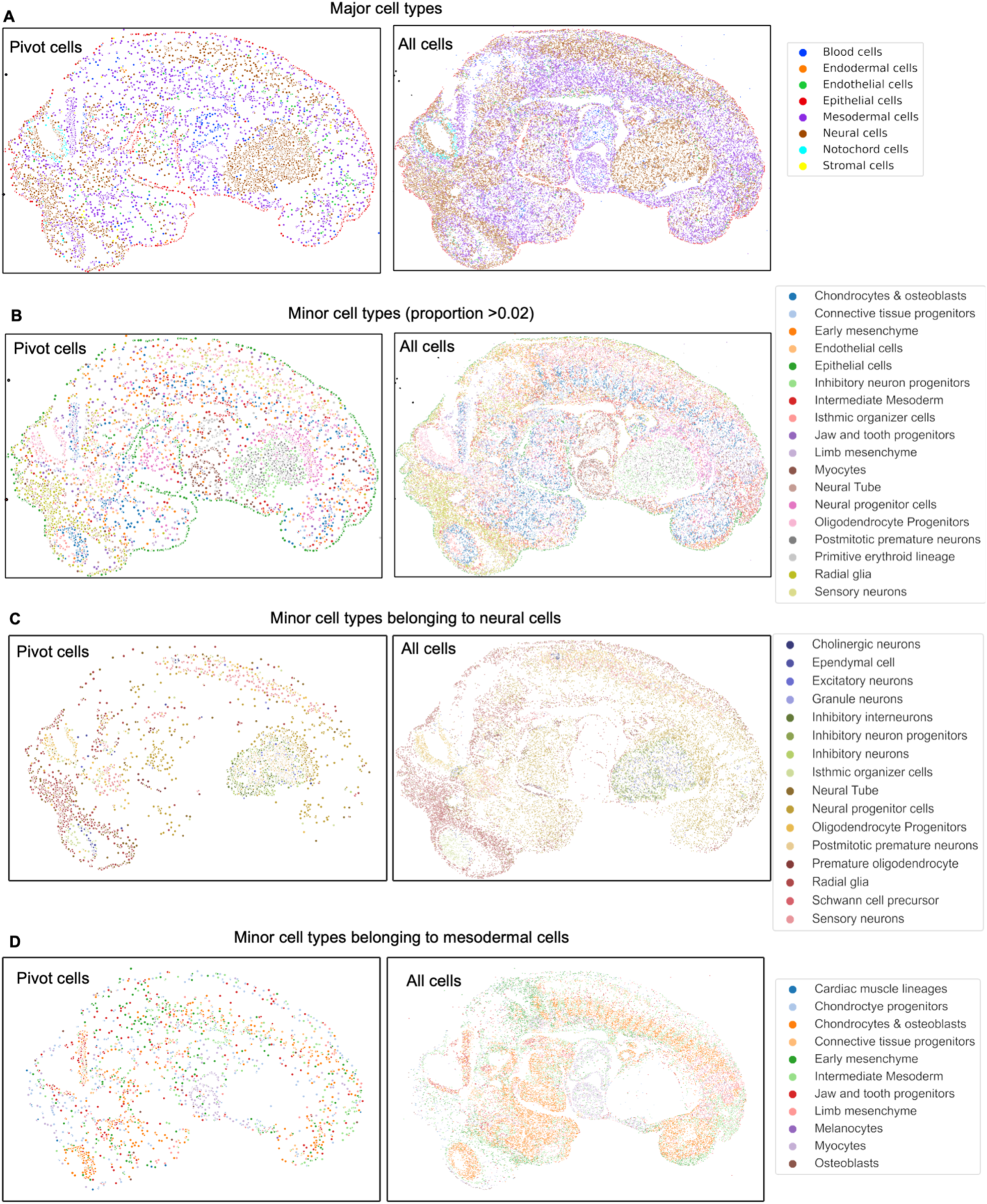
Deconvolution and WNN integration of DBiTplus datasets. **a** Top: UMAPs showing CODEX-informed deconvolution of DBiTplus feature only (Left) and a combination of DBiTplus and CODEX features (Right). Epithelial cells are better delineated when both deconvoluted DBiTplus and CODEX features are used via WNN integration. Bottom: No prior deconvolution with DBiTplus feature only (Left) and DBiTplus and CODEX feature (Right). The comparison with the top row demonstrates CODEX staining on the same tissue section markedly enhances the amount of information one gets from DBiTplus data. **b** Seurat WNN integration of DBiTplus+ and CODEX datasets. Weight of CODEX modality in Seurat WNN analysis. Strongest weights observed for T cell subtypes and lowest observed for B cell subtypes. These contributions are influenced by the composition of the CODEX marker panel.

**Extended Data Fig. 9.**
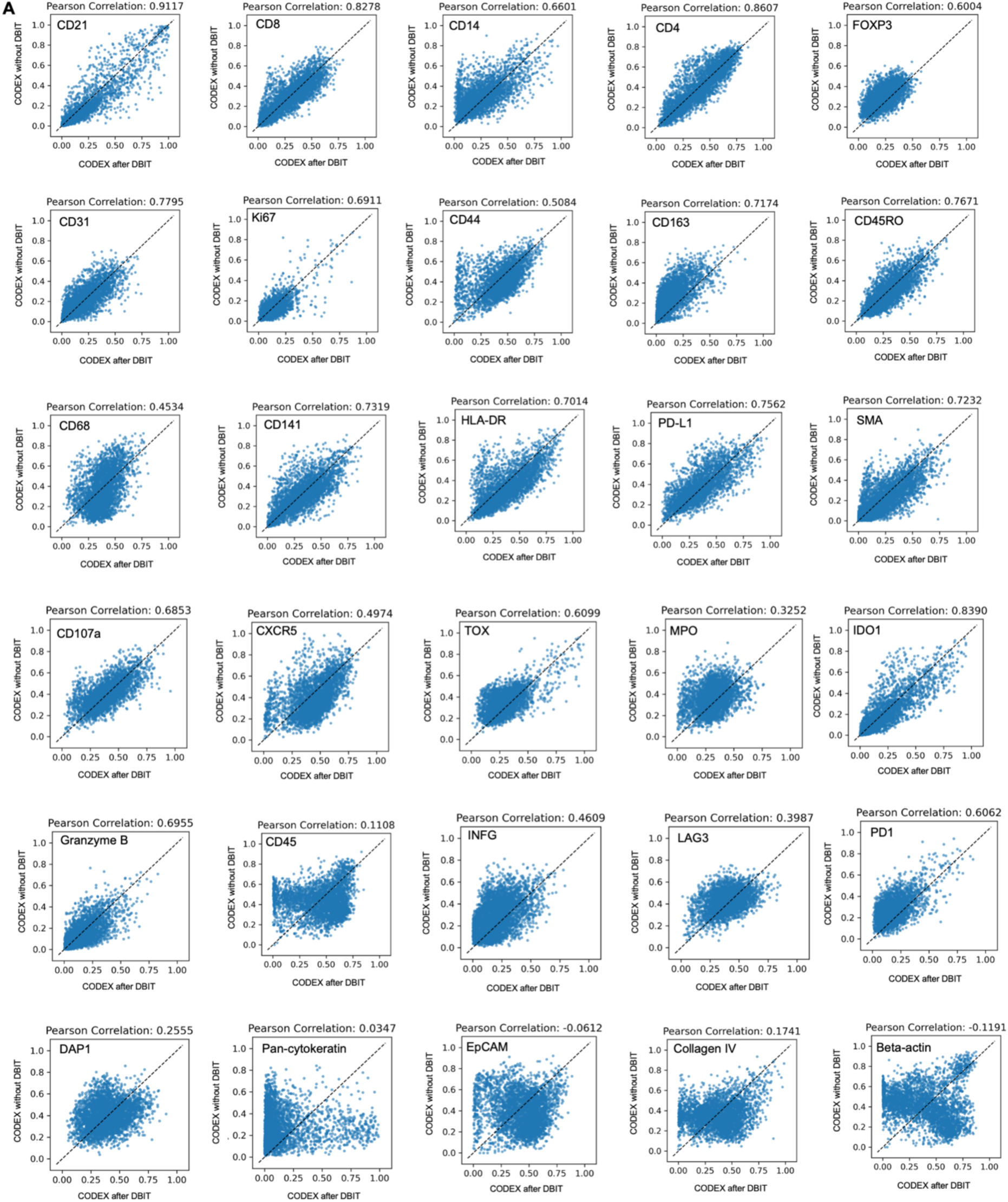
Assessing the quality of CODEX data from DBiTplus workflow. Correlation between tissue section stained via CODEX without prior DBiTplus and tissue section stained via CODEX after the DBiTplus workflow. Strong correlation is observed for most markers. Lowest correlations observed for markers such as Beta-actin and EpCAM.

**Extended Data Fig. 10.**
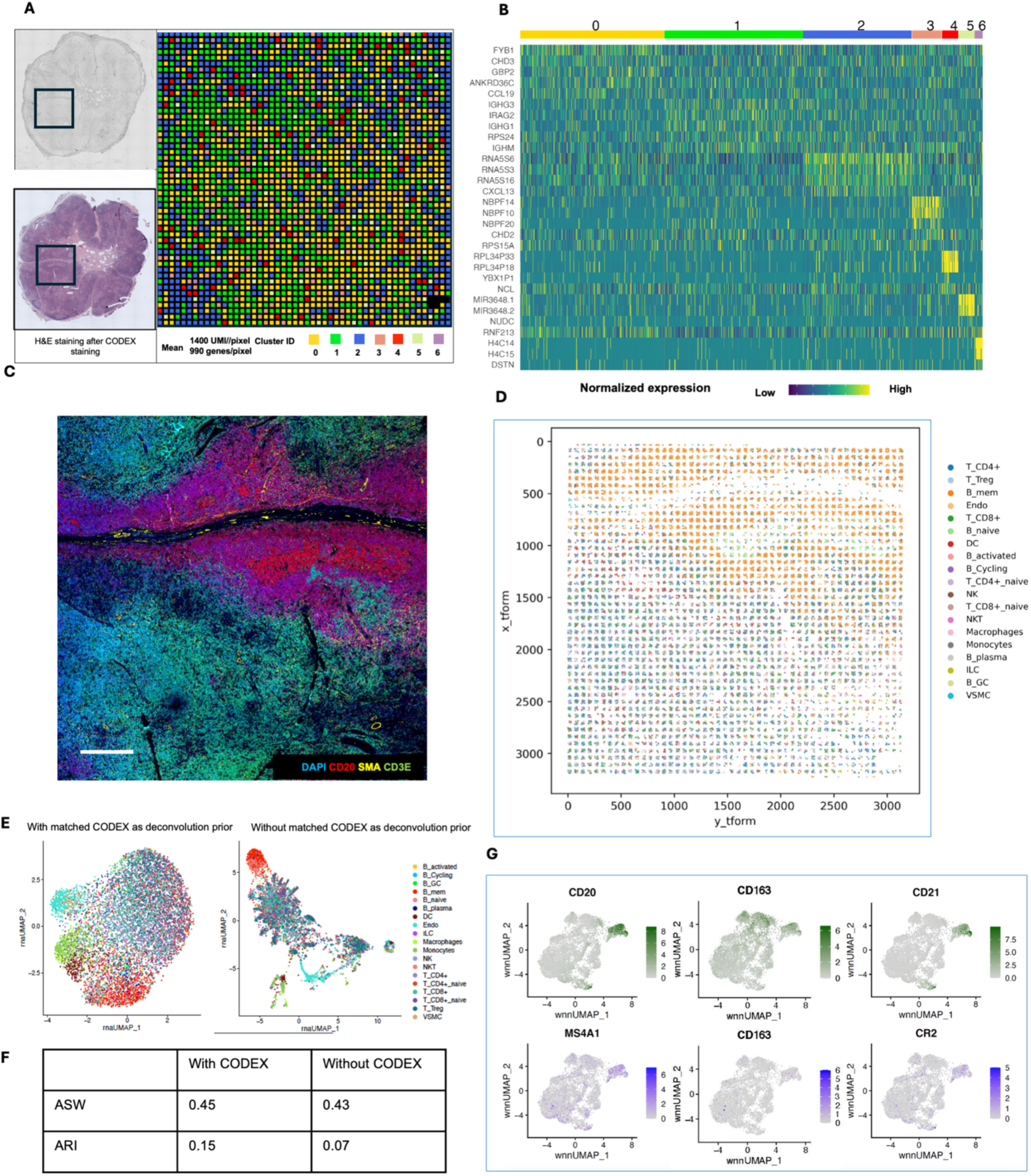
Spatial multi-omics profiling of FFPE human lymph node tissue section (25 μm resolution). **a** Spatial transcriptome distribution of FFPE human lymph node tissues barcoded using the microfluidic device with 25 μm resolution. Black box represents the DBiTplus ROI. Unsupervised clustering identified 7 distinct clusters whose spatial distribution closely matches the histological organization of human lymph nodes. **B** Heatmap showing the top 7 differentially expressed genes for each spatial cluster. **c** CODEX staining (panel of 35 markers) is performed on the same tissue section following the DBiTplus workflow. Scale bar 1mm. **d** Deconvolution of DBiTplus spots mapped by CODEX dataset. The identities of each cell within the DBiTplus spots can be directly inferred. **e** UMAPs of deconvoluted DBiTplus data: CODEX-informed deconvolution of DBiTplus feature (Left) and DBiTplus deconvolution without CODEX information (Right). **f** Higher ASW and ARI scores achieved by CODEX-informed deconvolution of DBiTplus spots suggesting better spot deconvolution quality. **g** Comparison of expression levels of protein features from CODEX (top) and corresponding RNA from DBiTplus after deconvolution (bottom).

**Extended Data Fig. 11.**
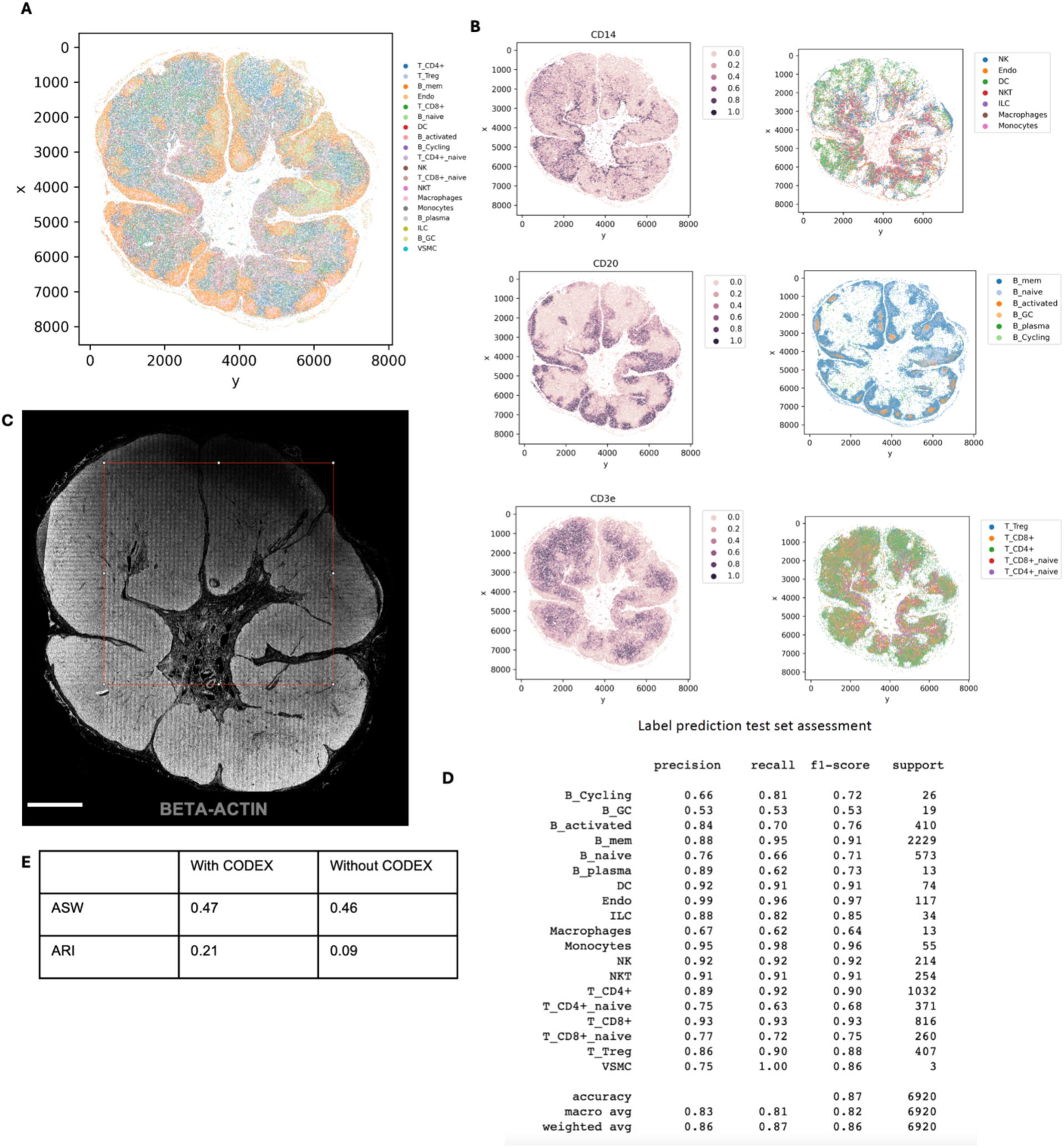
Spatial multi-omics profiling of FFPE of benign human lymph node tissue section (25 μm resolution). **A** Cell type annotation of whole CODEX dataset following integration with single cell RNA-seq reference dataset using MaxFuse. **b** Left: Spatial plots of different protein markers CD14 (Macrophages), CD20 (B cells) and CD3ɛ (T cells). Right: Plots of subtypes of the major cell types from the plots on the left. **c** Imprint of DBiTplus microfluidic channel on the tissue section observed in the Beta-actin channel. Scale bar 1mm**. d** Assessment of the label prediction test set. Highest f1-scores observed for endothelial cells, macrophages, dendritic cells and B cells. **e** Higher ASW and ARI scores for matched CODEX as deconvolution prior.

**Extended Data Fig. 12.**
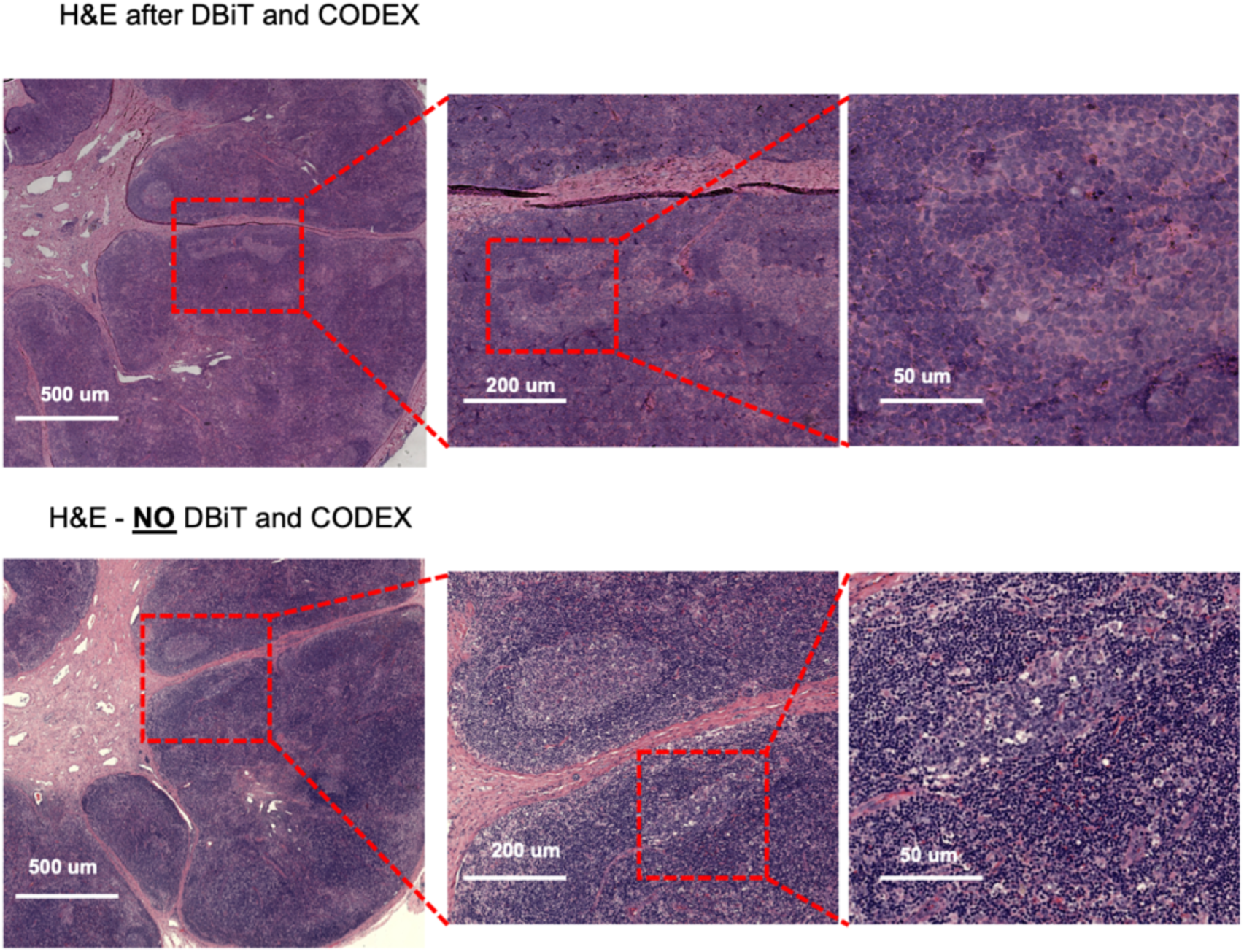
Comparison of H&E staining from FFPE of benign Human Lymph node sample after DBiTplus and CODEX workflows and an adjacent section without DBiTplus or CODEX. Identical regions from the adjacent sections undergo H&E staining. Comparable quality H&E is observed in both workflows.

**Extended Data Fig. 13.**
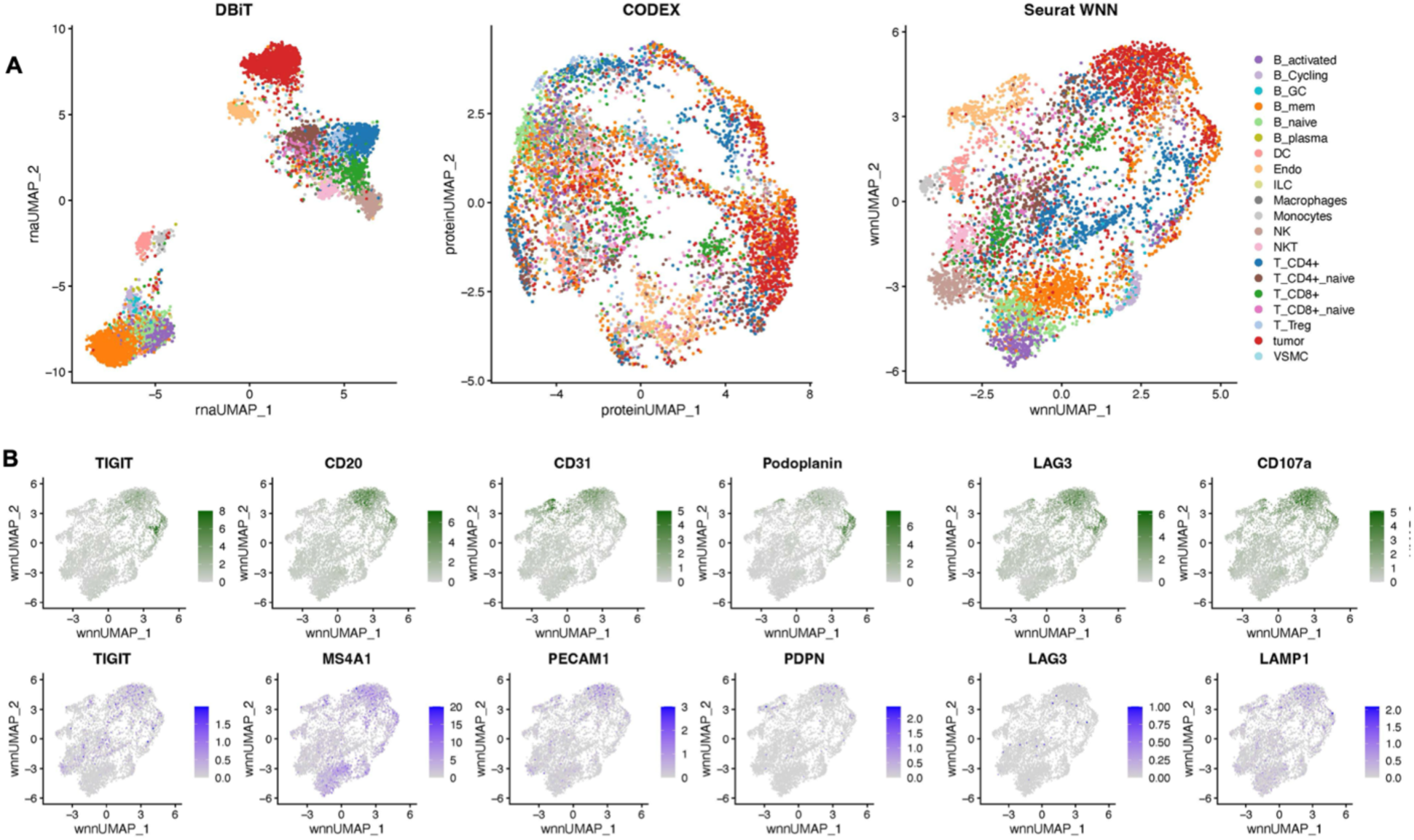
Integration of CODEX dataset and single cell RNA-seq reference dataset for DLBCL sample using MaxFuse. **(A)** UMAP of the integration of the DBiTplus and CODEX dataset showing distinct cell types. Tumor cells were first labeled by codex markers only. MaxFuse then used for annotating non-tumor cells. **(B)** Comparison of expression levels of protein features from CODEX (top) and corresponding RNA from DBiTplus after deconvolution (bottom).

